# Heterogeneity in inhibition of genetically diverse dengue virus strains by *Wolbachia*

**DOI:** 10.1101/2025.09.18.677129

**Authors:** Afeez Sodeinde, Emilie Finch, Ke Li, Mallery I. Breban, Ellie Bourgikos, Braiya L. Nolan, Nicole M. Feriancek, Isabel M. Ott, Julian W. Bakker, Verity Hill, Perran A. Ross, Xinyue Gu, Angkana T. Huang, Henrik Salje, Nathan D. Grubaugh, Chantal B.F. Vogels

## Abstract

The release of *Aedes aegypti* mosquitoes transinfected with the virus-inhibiting *Wolbachia* bacterium has the potential to reduce the burden caused by dengue virus (DENV). However, the robustness of this control strategy across the wide genetic diversity of DENV remains unknown. Here, we systematically tested two commonly used *Wolbachia* strains (*w*AlbB and *w*MelM) for their ability to inhibit 60 genetically diverse DENV isolates spanning all four serotypes. We found stronger inhibition by *w*MelM (median relative dissemination: 0.04) than *w*AlbB (median relative dissemination: 0.19). Furthermore, while we found substantial heterogeneity in inhibition across DENV isolates, we found that more DENV-3 isolates were weakly inhibited (median relative dissemination: 0.47 for *w*AlbB and 0.39 for *w*MelM) compared to the other serotypes (median relative disseminations: 0.10-0.18 for *w*AlbB and 0-0.11 for *w*MelM). Using transmission dynamic models, we further showed that differential *Wolbachia* inhibition results in increased probability of reemergence, particularly in high transmission intensity settings, with strong selection for DENV strains that have higher relative dissemination in mosquitoes. Our work highlights the importance of considering DENV genetic diversity, including the long-term risk of selection, in *Wolbachia*-based control interventions.

## Introduction

Nearly half of the world’s population is at risk of contracting infection with the mosquito-borne dengue virus (DENV)^1^. This virus is primarily transmitted by *Aedes aegypti* mosquitoes, and causes the greatest burden in South-East Asia and South America due to the tropical and subtropical climates being the most favorable environments for the survival of *Ae. aegypti* mosquitoes^2^. In the past five years, there has been a steady increase in reported dengue cases with a record number of 14.1 million cases reported in 2024, and substantial increases in the Americas^3,4^. The global distribution of dengue risk is also expected to continue to expand due to changing environmental suitability for mosquito vectors and local DENV transmission^5–7^. This increasing burden highlights the importance of developing sustainable prevention and control strategies that will be globally effective.

One promising approach for controlling DENV involves releasing *Ae. aegypti* mosquitoes transinfected with the endosymbiotic *Wolbachia pipientis* bacterium^8^. The *Wolbachia* release strategies aim to either suppress the local mosquito populations or replace them with *Wolbachia*-transinfected mosquitoes that have significantly reduced ability to transmit mosquito-borne viruses, including DENV^9,10,11^. Previous experimental and field studies have shown that variants of both the *Wolbachia w*Mel (derived from *Drosophila melanogaster*) and *w*AlbB (derived from *Ae. albopictus*) strains are effective in inhibiting the transmission of DENV by mosquitoes, resulting in mitigation of local transmission^10,12^. However, the underlying mechanisms of *Wolbachia* inhibition are not completely understood. Furthermore, *Wolbachia’s* inhibitory effect on DENV tends to be incomplete, resulting in occasional transmission. This raises a concern that virus selection may be a challenge to the long-term sustainability of this control strategy^13,14^.

One potential threat to the long-term effectiveness and sustainability of *Wolbachia*-based control is the high genetic diversity between and within the four DENV serotypes. DENV consists of four antigenically distinct serotypes (DENV-1 to DENV-4) that share approximately 65% genome similarity^15^. The serotypes are further divided into 4–7 genotypes, each with 6–8% genomic divergence^16^. Previous experimental studies showing DENV inhibition by *Wolbachia* have typically focused on a few selected virus isolates per serotype^17–20^. No study has comprehensively investigated *Wolbachia* genotype by DENV genotype interactions that included both the *w*Mel and *w*AlbB strains as well as the genotype diversity within the four DENV serotypes. We hypothesize that there is substantial variability in *Wolbachia* inhibition across genetically distinct DENV genotypes and *Wolbachia* strains. It is crucial to consider DENV genetic diversity in these studies as variation in inhibition may impact the outcomes of *Wolbachia* trials in the field.

This study addresses a critical knowledge gap in the understanding of the long-term implications of *Wolbachia* release on DENV diversity and epidemiology. Although mass release programs consider several factors to maximize introgression, successful introgression of *Wolbachia* strains that do not effectively block transmission could be highly problematic, as once a *Wolbachia* strain is established it is difficult to eradicate^21^. By systematically considering the wide diversity of DENV, we can understand the future population impacts of these release programs and better inform future control efforts. Our aim was to investigate the heterogeneity in virus inhibition between *Wolbachia* strains and a panel of genetically diverse DENV isolates. Overall, we found that inhibition was stronger by *w*MelM than *w*AlbB, with substantial variation in the level of inhibition of each of the DENV serotypes and isolates. Moreover, when modeling the impact of differential *Wolbachia* inhibition on DENV strain diversity, we found evidence for selection of DENV strains with higher relative dissemination over time under both *w*AlbB and *w*MelM scenarios. Our work highlights the importance of considering DENV genetic diversity in *Wolbachia*-based control interventions.

## Results

### Establishing an experimental system to study dengue virus by *Wolbachia* interactions

We developed a comprehensive experimental system encompassing three *Ae. aegypti* mosquito colonies and a panel of genetically diverse DENV isolates to robustly evaluate *Wolbachia* inhibition on virus infection and dissemination. Our mosquito colonies include wildtype *Ae. aegypti* (not transinfected with *Wolbachia*; WT), and *Ae. aegypti* from the same genetic background that were stably transinfected with *w*AlbB or *w*MelM (see Methods). We exposed these three mosquito colonies to genetically diverse DENV isolates from collections provided by the Yale Arbovirus Research Unit and World Reference Center for Emerging Viruses and Arboviruses. Our final panel included twenty isolates each of DENV-1 and DENV-2, eleven isolates of DENV-3, and nine isolates of DENV-4 (**Fig. 1a; Supplementary Table 1**). All major genotypes of DENV-1 (1I-1V and VII) and DENV-2 (2I to 2VI) were represented by at least one DENV isolate, and three out of four genotypes of DENV-3 (3I to 3III) and DENV-4 (4I, 4II, 4IV) were represented in the panel, due to availability of virus isolates from both collections (**Fig. 1a-b**). Our panel of DENV isolates originated from 25 countries in Asia, Africa, and the Americas, span a 72-year collection period (1944 to 2016), and were isolated from humans (n=46), mosquitoes (n=5) and a Southern pig-tailed macaque (n=1; **Supplementary Table 1**). By testing our panel of 60 DENV isolates in three *Ae. aegypti* colonies (WT, *w*AlbB, *w*MelM) resulting in 180 unique mosquito by DENV combinations, we established a comprehensive study system to determine the impact of DENV genetic diversity on inhibition by *Wolbachia*.

**Fig. 1:**
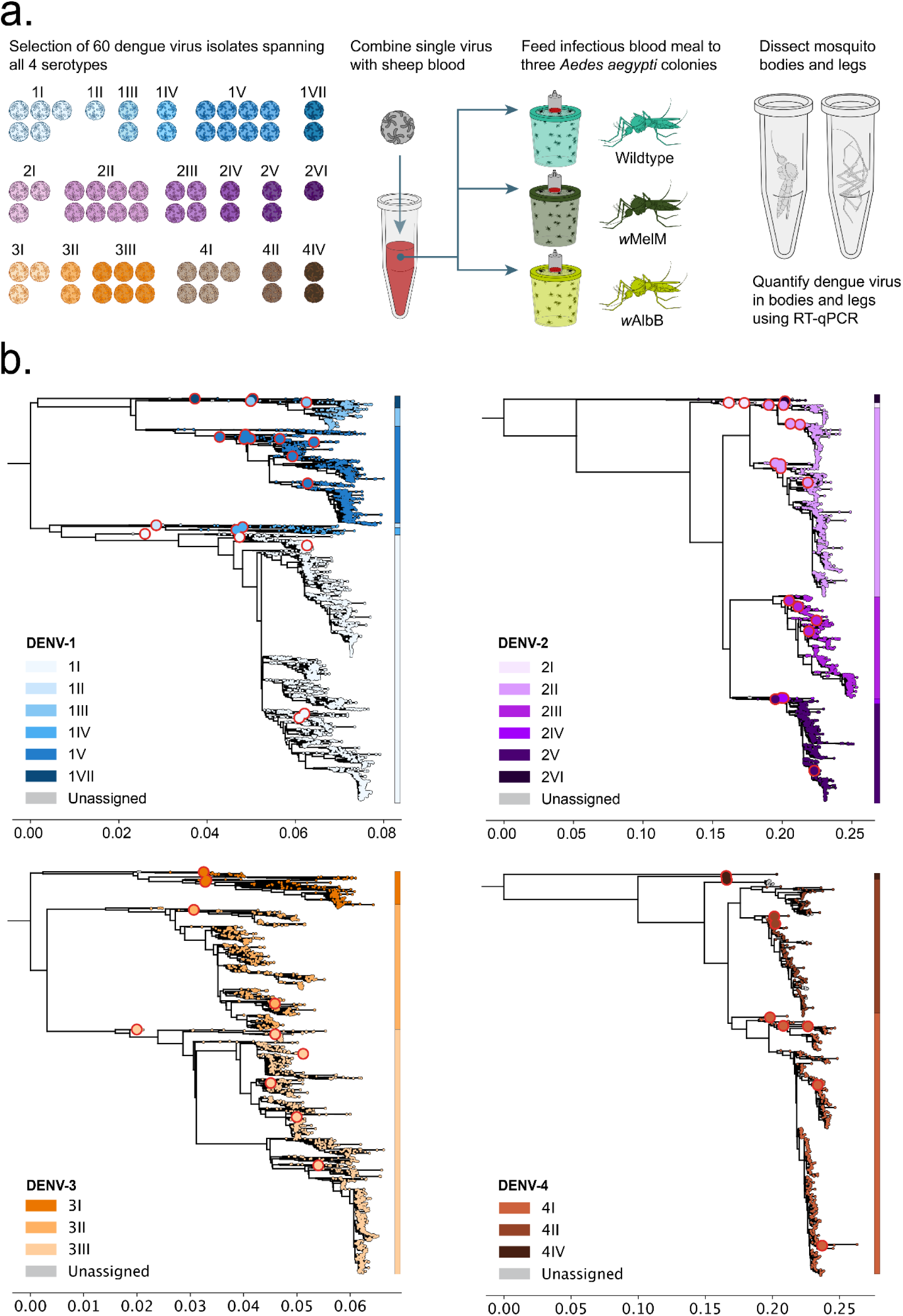
Overview of experimental study system to determine the impact of DENV genetic diversity on inhibition by *Wolbachia*. a, Overview of experimental design with a panel of 60 genetically diverse DENV isolates tested in three *Aedes aegypti* colonies (wildtype; not transinfected with *Wolbachia* [WT], and two strains of *Wolbachia* [*w*AlbB and *w*MelM]). Each DENV isolate was harvested on the day of mosquito infections, and used to prepare three Hemotek feeders to allow parallel feeding of the three mosquito colonies on the same virus. Engorged female mosquitoes were incubated for 10 days and mosquito bodies and legs were tested for DENV infection and dissemination, respectively, using RT-qPCR. b, Maximum likelihood phylogenetic trees were generated using a subset of publicly available DENV genomes to show genetic diversity within the major genotypes of each serotype, stratified by serotype, and colored by genotype. The red outlined circles denote the placement of DENV isolates selected for this study, highlighting their genetic diversity amongst the known global virus population.

### Variation in infection and dissemination between mosquito colonies and DENV isolates

To determine the impact of DENV genetic diversity on inhibition by *Wolbachia*, we exposed three *Ae. aegypti* colonies (WT, *w*AlbB, *w*MelM) in parallel to our panel of 60 genetically diverse DENV isolates (**Fig. 1a**). After 10 days of incubation, we determined infection (presence of DENV in mosquito body; **Fig. 2a**) and dissemination rates (presence of DENV in mosquito legs; **Fig. 2b**) using RT-qPCR. Our sample size across all 180 unique mosquito-virus combinations (3 colonies x 60 DENV isolates) included 5,283 exposed mosquitoes that were tested for DENV infection, with an average sample size of 29 mosquitoes per experimental condition. Overall, we observed higher infection rates as compared to dissemination rates, consistent with the known role of the mosquito midgut as a barrier to dissemination^22,23^.

**Fig. 2:**
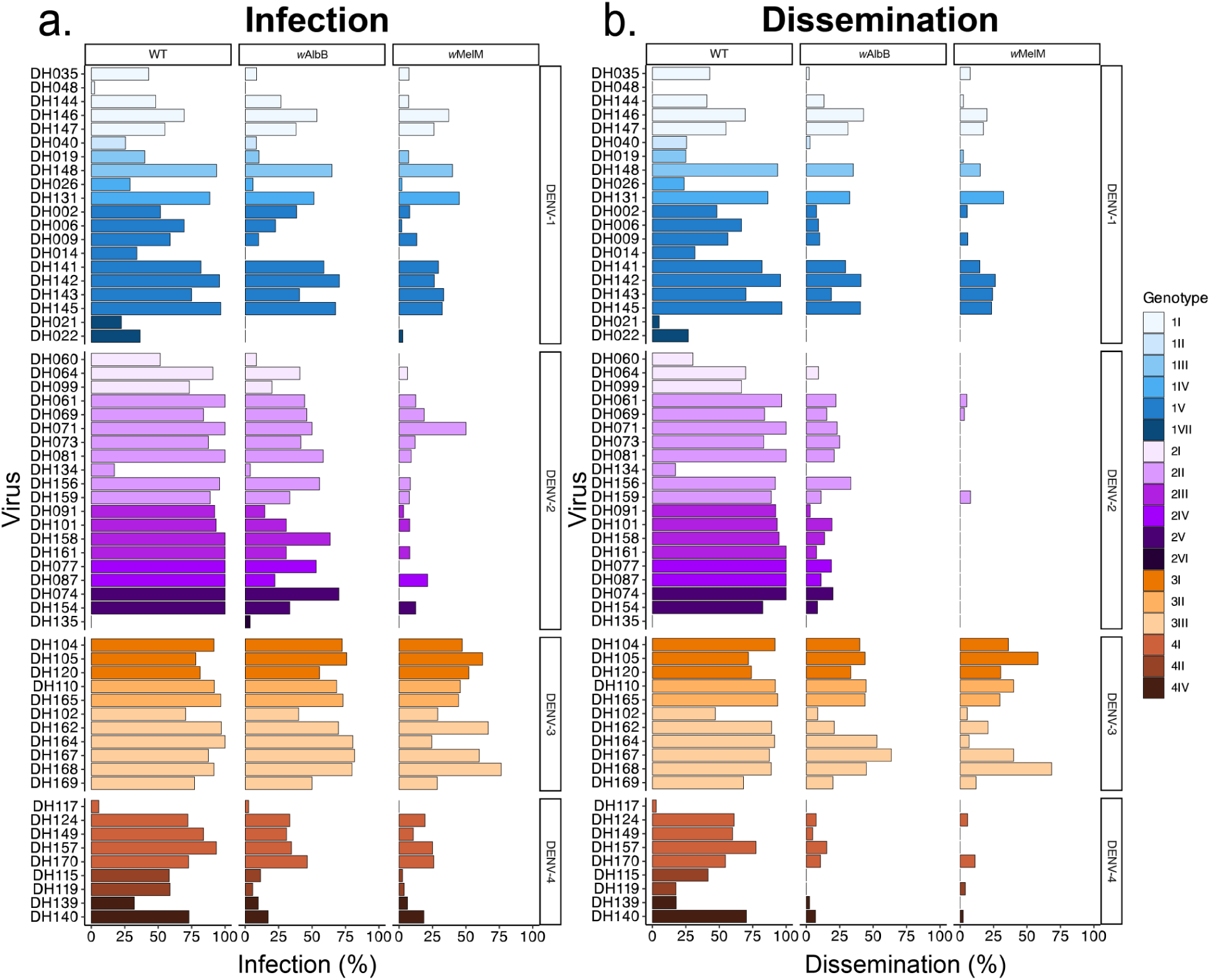
Infection and dissemination rates vary by *Ae. aegypti* colony (presence/absence of *Wolbachia*), *Wolbachia* strain, and DENV isolate. Dengue virus infection and dissemination rates were determined 10 days after exposing WT, *w*AlbB, and *w*MelM mosquitoes to an infectious blood meal (all three colonies exposed to the same virus stock in parallel). a, Infection rates of each DENV isolate (rows) for each of the three *Ae. aegypti* colonies (columns). b, Dissemination rates of each DENV isolate (rows) for each of the three *Ae. aegypti* colonies (columns). Infection and dissemination rates were calculated as the percentage of positive mosquito bodies and legs, respectively, out of the total exposed mosquitoes. DENV isolates are colored by serotype and genotypes with DENV-1 = blue, DENV-2 = purple, DENV-3 = orange, DENV4 = brown. Metadata for each of the DENV isolates is included in **Supplementary Table 1**. Underlying data, including sample sizes, for this figure are available in the **Source Data File**.

As dissemination has been shown to be a reliable proxy for transmission potential^24^, we further compared dissemination rates between mosquito colonies and DENV isolates. First, we compared dissemination rates in WT mosquitoes, which were expected to be the most susceptible to DENV. We found a high level of variability in dissemination rates ranging from 0-100% (DENV-1: 0-97%, DENV-2: 0-100%, DENV-3: 47-94%, and DENV-4: 3-77%) even in absence of *Wolbachia* (**Fig. 2b**). This supports prior studies indicating that vector competence, the ability of a mosquito to become infected and eventually transmit a virus, is dependent on the specific combination of mosquito and virus genetic background^25^.

Next, we determined dissemination rates of our panel of DENV isolates in *w*AlbB and *w*MelM *Wolbachia*-transinfected colonies. While previous studies revealed *Wolbachia* strain-specific inhibition of DENV, with intermediate DENV inhibition by *w*AlbB and strong inhibition by *w*MelM^26,27^, we found a high level of variability in inhibition by both *Wolbachia* strain and DENV isolate. Dissemination rates ranged between 0-64% (DENV-1: 0-43%, DENV-2: 0-33%, DENV-3: 9-64%, and DENV-4: 0-15%) in *w*AlbB transinfected mosquitoes and between 0-68% (DENV-1: 0-33%, DENV-2: 0-8%, DENV-3: 5-68%, and DENV-4: 0-11%) in *w*MelM transinfected mosquitoes (**Fig. 2b**). Our findings reveal high variability in infection and dissemination by mosquito colony (presence/absence of *Wolbachia*), *Wolbachia* strain, and DENV isolate.

### *w*MelM shows stronger inhibition than *w*AlbB, with some DENV-3 isolates being less inhibited by both *Wolbachia* strains

The high variation in infection and dissemination of the diverse DENV isolates that we observed in WT mosquitoes makes it hard to directly compare the level of inhibition between *Wolbachia* strains and DENV serotypes. To account for this variation, we calculated DENV infection (**Supplementary Fig. 1**) and dissemination (**Fig. 3**) in *w*AlbB and *w*MelM relative to WT mosquitoes, with a value of 0 meaning no infection/dissemination (complete inhibition) and a value of 1 meaning infection/dissemination equal to WT (no inhibition).

**Fig. 3:**
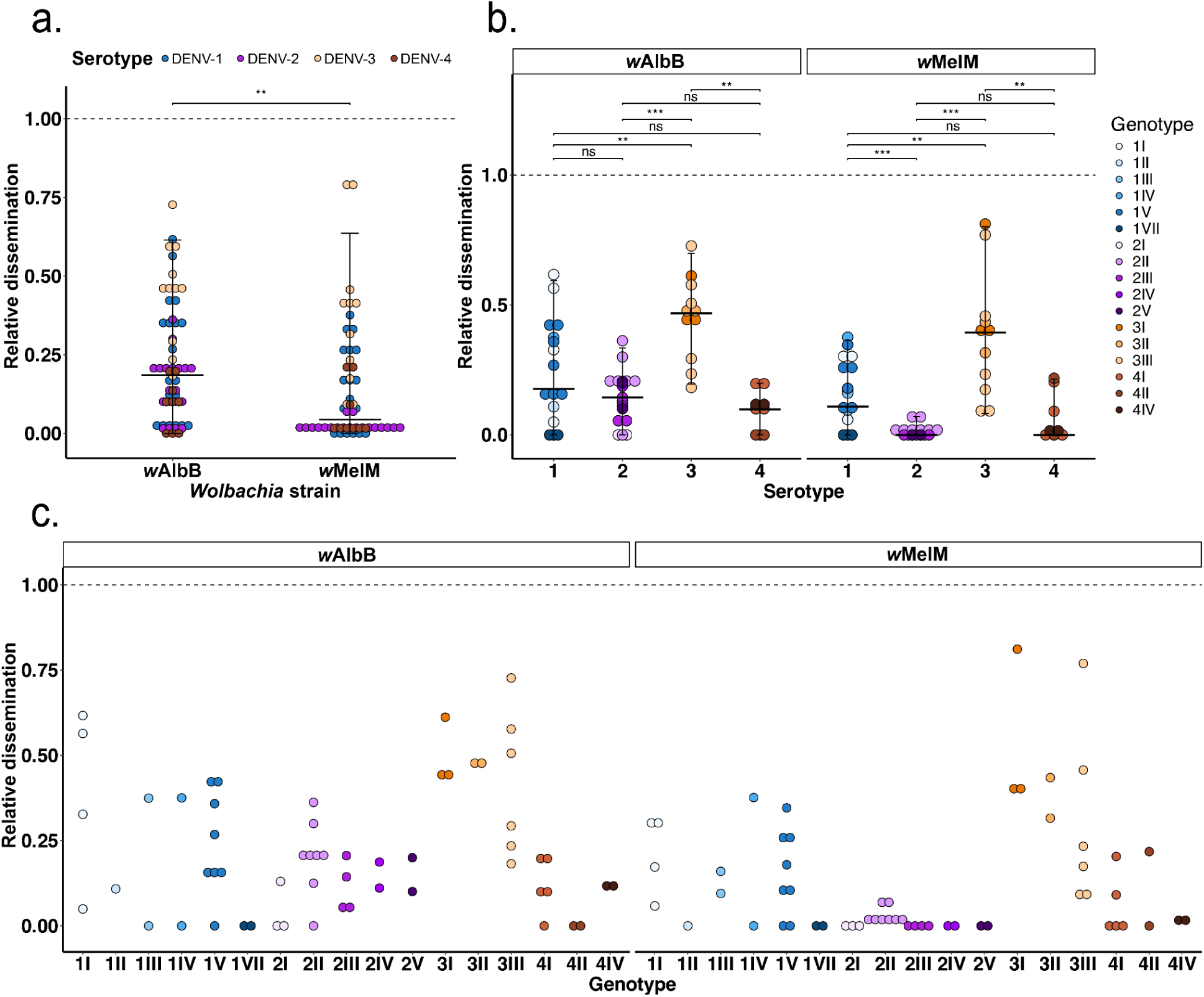
*Wolbachia* strain and DENV serotype significantly impact the relative dissemination of DENV. Relative dissemination for each DENV isolate was calculated as the ratio of dissemination rate in *w*AlbB or *w*MelM mosquitoes relative to dissemination rate in WT mosquitoes. a, Relative dissemination rate of *w*AlbB and *w*MelM mosquitoes for all virus isolates. b, DENV serotype-level relative dissemination rates for *w*AlbB and *w*MelM mosquitoes. c, DENV genotype-level relative dissemination rates for *w*AlbB and *w*MelM mosquitoes. For figures a-c, each dot represents the relative dissemination of a DENV isolate in either *w*AlbB or *w*MelM mosquitoes relative to WT, with 0 = no dissemination relative to WT (complete inhibition) and 1 = equal dissemination relative to WT (no inhibition). Dots are colored by DENV serotype, with DENV-1 = blue, DENV-2 = purple, DENV-3 = orange, DENV4 = brown. Horizontal lines indicate the median with 95% quantile intervals, corresponding to the range between the 2.5th and 97.5th percentiles. The Wilcoxon test was used for statistical comparisons with Benjamini-Hochberg correction to account for multiple comparisons. In all panels, *** = p < 0.001, ** = p < 0.01, * = p < 0.05 and ns ≥ 0.05. Underlying data for this figure are available in the **Source Data File**.

We compared relative dissemination between *w*AlbB and *w*MelM to identify *Wolbachia* strain-specific effects on DENV inhibition. Our analysis showed that *w*MelM (median relative dissemination: 0.04 [95% quantile interval: 0-0.64]) was overall more effective at inhibiting DENV dissemination when compared to *w*AlbB (median relative dissemination: 0.19 [95% quantile interval: 0-0.62]; **Fig. 3a**). However, we observed substantial variation in inhibition within both *w*MelM and *w*AlbB colonies and hypothesized that DENV serotype- or genotype-specific effects may cause the observed variation. Thus, we tested the impact of DENV serotypes on relative dissemination by each *Wolbachia* strain. While we found substantial variation among the DENV isolates, we found that many DENV-3 isolates were minimally inhibited by both *Wolbachia* strains (median relative dissemination: 0.47 [95% quantile interval: 0.19-0.70] for *w*AlbB and 0.39 [95% quantile interval: 0.08-0.80] for *w*MelM) as compared to isolates from the other three serotypes (median relative disseminations: 0.10-0.18 for *w*AlbB and 0-0.11 for *w*MelM; **Fig. 3b**). Moreover, many DENV-2 isolates had high levels of inhibition by *w*MelM (median relative disseminations range: 0 [95% quantile interval: 0-0.07]), whereas this effect was not seen in the inhibition of DENV-2 by *w*AlbB (median relative disseminations range: 0.14 [95% quantile interval: 0-0.33]). This *Wolbachia*-specific effect on DENV-2 was particularly clear when we examined the correlation between relative dissemination in *w*MelM compared to *w*AlbB. We found a moderate to strong correlation between *w*MelM and *w*AlbB inhibition for DENV-1 and DENV-3, but not for DENV-2 and DENV-4 (**Supplementary Fig. 2**). Analyzing the serotype-specific effects for each *Wolbachia* strain also revealed that relative dissemination rates varied widely within each of the DENV serotypes. A further comparison at the DENV genotype and sub-genotype level showed no clear pattern of inhibition, suggesting that even at these levels, there may be other confounding factors that could impact the variability of *Wolbachia* inhibition (**Fig. 3c**). In summary, our findings reveal a high level of heterogeneity in relative DENV dissemination, with specific interactions between *Wolbachia* strain and DENV isolate determining the level of inhibition.

### Input stock virus titers impact DENV dissemination in a serotype-specific manner

To avoid reduced infectivity due to freezing DENV stocks^28,29^, we immediately used DENV stocks harvested from cell culture to prepare the infectious blood meals, resulting in variation in input stock virus titers. To further investigate the impact of variation in input stock titers on DENV dissemination, we performed a correlation test between input stock viral titers and dissemination rates for all tested combinations (**Fig. 4)**. We observed a significant and strong positive correlation between input stock titers and dissemination of DENV-1 and DENV-2 viruses in WT mosquitoes. This confirms that a higher infectious dose generally results in higher infection and dissemination as observed by previous studies (**Fig. 4**; **Supplementary Fig. 3**)^30^. Compared to WT mosquitoes exposed to DENV-1 viruses, *w*AlbB mosquitoes had a slightly weaker correlation, followed by *w*MelM, which had the weakest correlation of all three colonies with DENV-1 input stock titers. This suggests that both *Wolbachia* strains were able to partially inhibit DENV-1, even with high input stock virus titers. Despite a significant and strong positive correlation between DENV-2 input stock titers and dissemination rates in WT, there was no significant correlation seen in either the *w*AlbB or *w*MelM colonies. This shows that both *Wolbachia* strains were able to inhibit DENV-2 across a range of virus titers in the infectious blood meal. In contrast, we found no correlation between input stock virus titers and dissemination rates for any of the mosquito colonies infected with DENV-3 and DENV-4 viruses. Overall, our findings show that input stock titers impact mosquito infection and dissemination mostly in WT mosquitoes, and that the observed variation in *Wolbachia* inhibition cannot be fully explained by input DENV stock titer in the infectious blood meal.

**Fig. 4:**
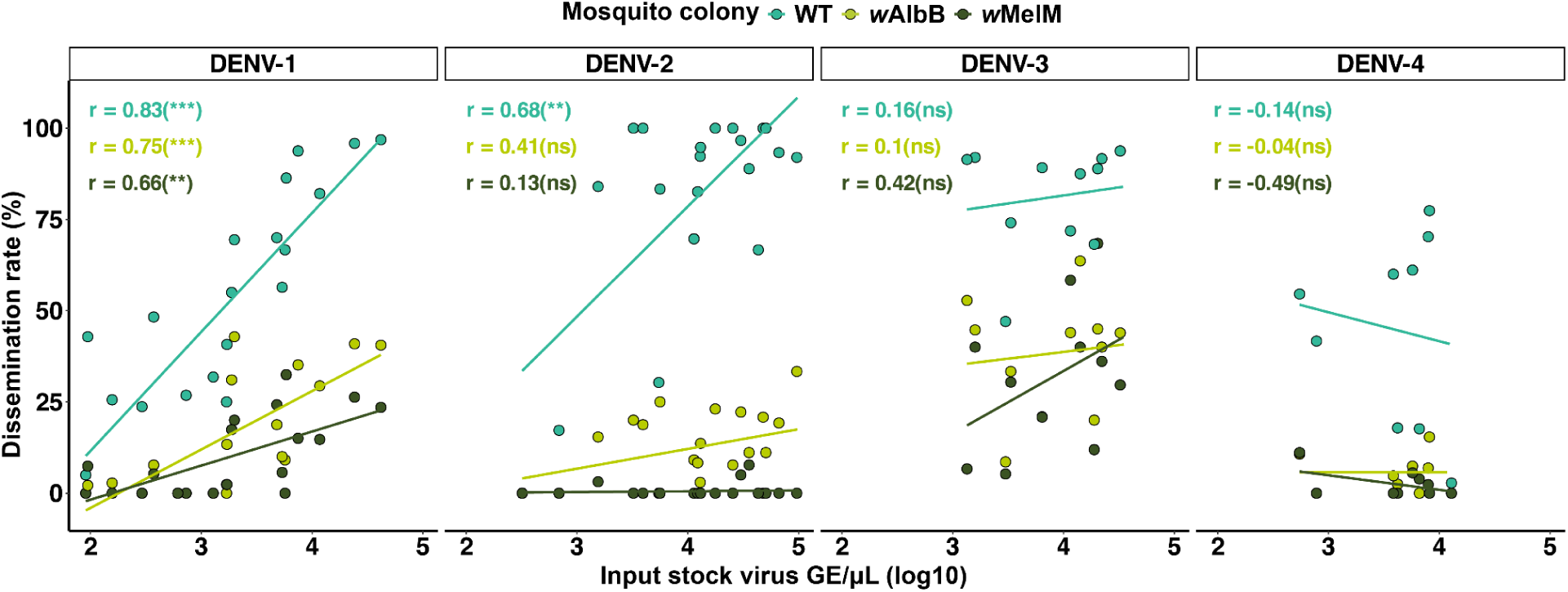
Impact of input stock virus titer in the infectious blood meal on dissemination rates across each DENV serotype. Correlation plots of input stock virus titers (measured as genome equivalents) with dissemination rates. Each dot represents a tested DENV isolate, and colored by mosquito colony with WT = teal, *w*AlbB = bright yellow-green, and *w*MelM = olive-green. In all panels, r = Pearson correlation coefficient, *** = p < 0.001, ** = p < 0.01, * = p < 0.05 and ns ≥ 0.05. Underlying data for this figure are available in the **Source Data File**.

### DENV inhibition and replication in mosquitoes is not impacted by *Wolbachia* density

One possible explanation for the lower infection and dissemination rates in the presence of *Wolbachia* is that DENV replication is impaired by *Wolbachia*. By quantifying DENV titers, measured as genome equivalents using RT-qPCR, in fully disseminated mosquitoes (mosquitoes that tested positive for DENV in both body and legs after the incubation period), we determined whether *Wolbachia* impacts DENV replication throughout the mosquito body. Our sample size of fully disseminated mosquitoes differed per mosquito colony ranging from highest for WT, intermediate for *w*AlbB, and lowest for *w*MelM, reflecting the different levels of inhibition induced by both *Wolbachia* strains (**Fig. 1b**; **Fig. 5a**). When comparing DENV titers between the three mosquito colonies, we found minor differences in DENV replication when stratified by serotype. We observed similar levels of replication in WT and *w*AlbB mosquitoes, except for DENV-2, where DENV titers were significantly higher in WT mosquitoes (median: 3.89 log_10_ GE/μL) as compared to *w*AlbB (median: 3.63 log_10_ GE/μL; **Fig. 5a**). DENV titers were similar between WT and *w*MelM mosquitoes, except for DENV-3, where *w*MelM titers (median: 3.13 log_10_ GE/μL) were significantly lower than WT (median: 3.48 log_10_ GE/μL; **Fig. 5a**). Moreover, we found similar DENV titers in *w*AlbB- and *w*MelM-transinfected mosquitoes, except for DENV-3, where DENV titers were significantly lower in *w*MelM (median: 3.13 log_10_ GE/μL) as compared to *w*AlbB (median: 3.45 log_10_ GE/μL; **Fig. 5a**). While we detected minor differences between some groups, there was significant overlap in the range of DENV levels in each of the three mosquito colonies. This suggests that all four DENV serotypes are capable of reaching similar levels of replication in presence or absence of *Wolbachia.* Together this indicates that, even though there is slight variation in DENV titers within and between colonies, *Wolbachia w*AlbB and *w*MelM did not significantly impact the ability of DENV to replicate to high levels in fully disseminated mosquitoes.

**Fig. 5:**
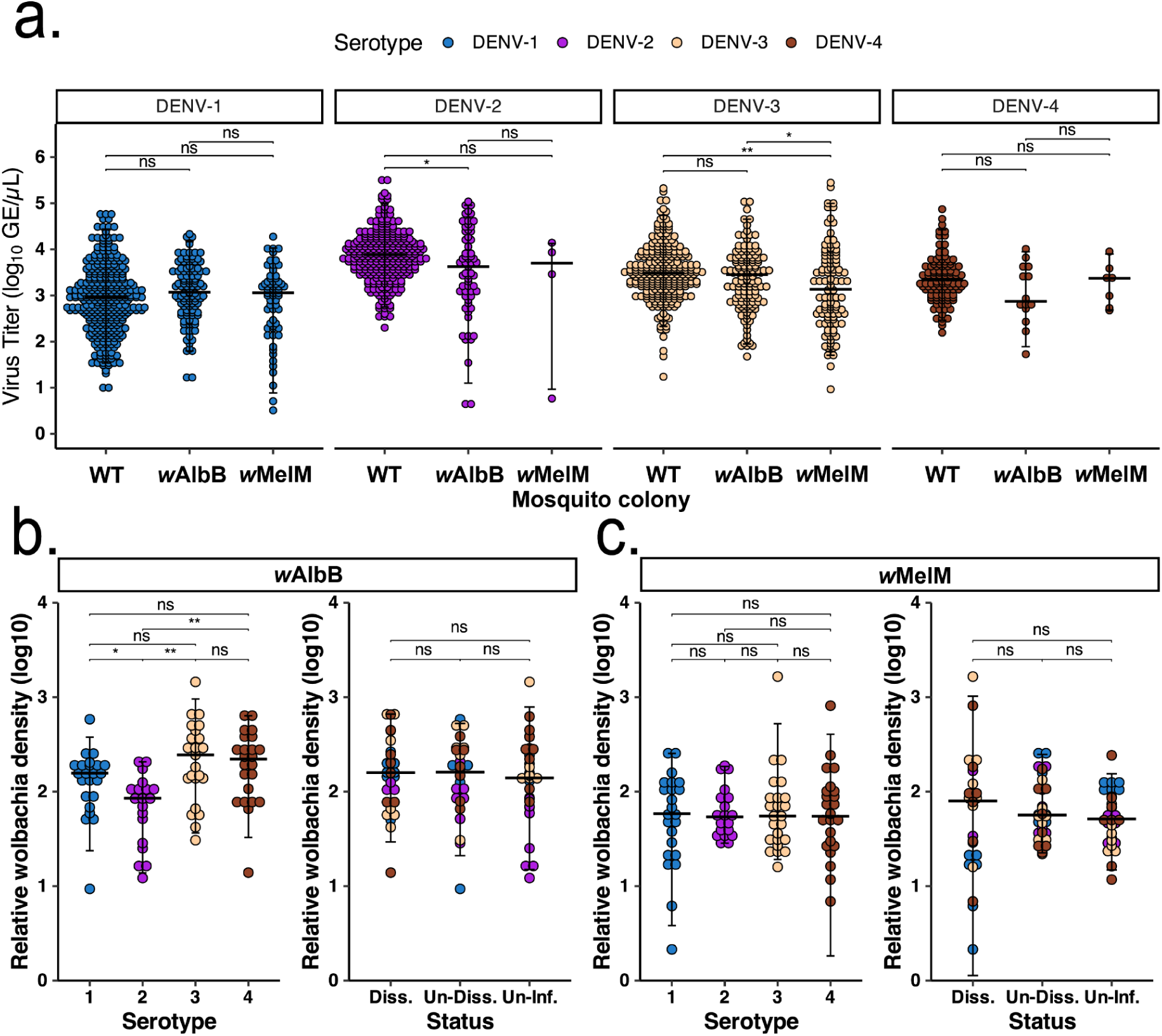
DENV inhibition and replication is not impacted by *Wolbachia* relative densities. a, DENV genome equivalents were quantified in the bodies of fully disseminated WT, *w*AlbB, and *w*MelM mosquitoes using RT-qPCR. Sample size was between 132-333 for WT, 23-129 for *w*AlbB and 4-96 for *w*MelM mosquitoes. b, Relative density of *Wolbachia w*AlbB strain determined for a random subset of mosquito bodies by DENV serotype (left) and dissemination status (right) using qPCR. Sample size was between 19-23 per serotype and 28-30 per dissemination status. c, Relative density of *Wolbachia w*MelM strain determined for a random subset of mosquito bodies by DENV serotype (left) and dissemination status (right) using qPCR. Sample size was between 19-24 per serotype and 28-32 per dissemination status. Each dot represents an individual mosquito colored by DENV serotype, with DENV-1 = blue, DENV-2 = purple, DENV-3 = orange, DENV4 = brown. Dissemination status is abbreviated as follows: Diss = Exposed mosquito with virus positive body and legs after incubation period, Un-Diss = Exposed mosquito with virus positive body and negative legs after incubation period, Un-Inf = Exposed mosquito with virus negative body and legs after incubation period. Horizontal bars indicate the median with 95% quantile intervals, corresponding to the range between the 2.5th and 97.5th percentiles. The Wilcoxon test was used for statistical comparison with Benjamini-Hochberg correction to account for multiple comparisons. *** = p < 0.001, ** = p < 0.01, * = p < 0.05 and ns ≥ 0.05. Underlying data for this figure are available in the **Source Data File**.

Given that the scale of our experiments required mosquito infections to be performed in a total of three batches, we wanted to rule out the possibility that unintended variability in relative *Wolbachia* density could impact observed heterogeneity in relative inhibition or levels of DENV replication. We, therefore, determined relative *Wolbachia w*AlbB and *w*MelM densities within randomly selected mosquitoes with varying virus titer and dissemination statuses (**Fig. 5b-c**). First, we compared relative *Wolbachia* densities in mosquitoes that were exposed to the four different DENV serotypes. For *w*AlbB, we found that relative densities were similar in mosquitoes exposed to DENV-1 (median: 2.20 log_10_), DENV-3 (median: 2.39 log_10_), and DENV-4 (median: 2.35 log_10_), whereas relative *Wolbachia* densities were significantly lower in mosquitoes exposed to DENV-2 (median: 1.93 log_10_) compared to the other three serotypes. When compared by infection and dissemination status, all exposed *w*AlbB mosquitoes had similar *Wolbachia* relative densities (median: 2.15-2.21 log_10_). We found no differences in relative *w*MelM densities when comparing mosquitoes exposed to any of the four DENV serotypes (median: 1.73-1.77 log_10_), or when comparing relative densities by dissemination status (median: 1.71-1.90 log_10_). These findings demonstrate that heterogeneity observed in DENV inhibition and high DENV replication in *Wolbachia*-transinfected mosquitoes cannot be explained by variation in relative *Wolbachia* densities in mosquito colonies.

### Mathematical modelling reveals the potential for *Wolbachia* mediated DENV strain selection

To explore the potential implications of the heterogeneity in *Wolbachia* inhibition, we used our experimental data to inform a stochastic transmission model (**Fig. 6**). First, we explored the impact of differing levels of relative dissemination across strains on DENV reemergence within the same population (**Fig. 6a**). We repeatedly simulated transmission dynamics considering four strains (representing one from each serotype) prior to *Wolbachia* introduction. Following introduction, three strains were completely inhibited while the inhibition level of the fourth strain varied systematically between 0 and 1. We found that the time to reemergence of the partially inhibited strain decreased with increasing levels of relative dissemination and increasing underlying R_0_ (**Fig. 6c**). With a low underlying transmission intensity of R_0_=1.5 we saw no reemergence within 75 years at relative dissemination levels at 0.55 or under and reemergence within 16 years (95% CrI: 12-19) at a high relative dissemination level of 0.8. At R_0_=4, we saw reemergence within our simulation period where relative dissemination was as low as 0.25, occurring within 4 years when relative dissemination was only 0.8 .

**Figure 6:**
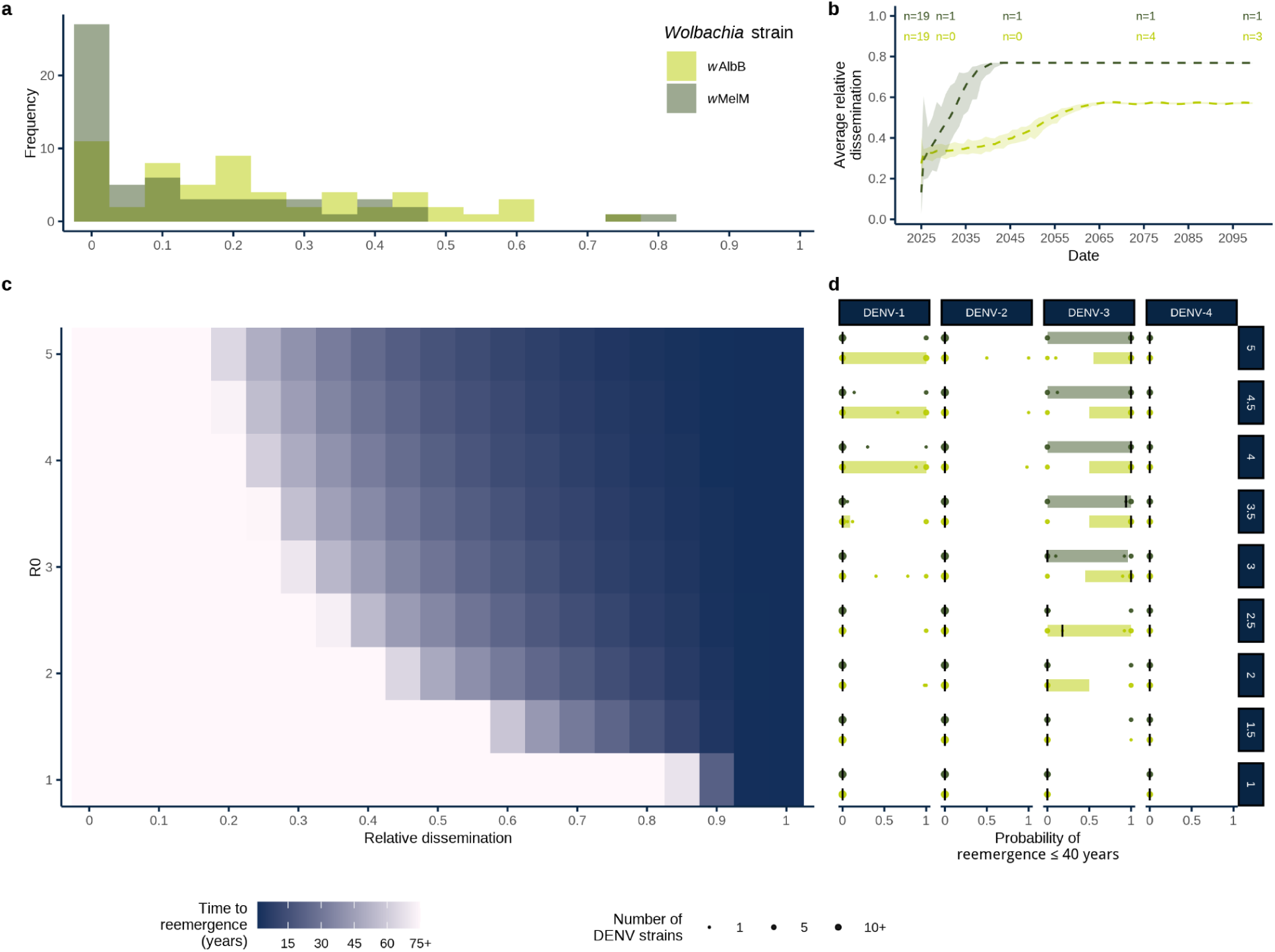
Potential impacts of differential *Wolbachia* inhibition on DENV reemergence and strain diversity. a, Histogram of relative dissemination levels of DENV strains in *w*AlbB (in bright yellow-green) and *w*MelM (in olive-green). b, Average relative dissemination of circulating DENV strains over time. The dashed line shows the median value over all 1000 stochastic simulations for *w*AlbB and *w*MelM (in bright yellow-green and olive-green, respectively) while the shaded area shows the 95% credible interval. The median number of circulating DENV strains is shown at the top of the panel for pre-release, 5 years post-release, 20 years post-release, 50 years post-release and 75 years post-release. Each simulation started with five DENV strains per serotype (20 strains total) under both *w*AlbB and *w*MelM interventions. By the end of the simulation period, we found a median of only two remaining circulating DENV strains under both *Wolbachia* scenarios. c, Estimated time to reemergence (in years) of DENV under different assumed transmission intensities (R_0_) and relative dissemination in *Wolbachia*-transinfected mosquitoes. Tiles where no reemergence occurred in the simulation period are marked as 75+ years. d, Probability of reemergence in ≤ 40 years for each DENV strain in the panel tested, shown separately by serotype. Black lines show the median probability of reemergence in ≤ 40 years by serotype, R_0_ and *Wolbachia* strain, shaded bars show the 95% credible interval in bright yellow-green and olive-green for *w*AlbB and *w*MelM, respectively. Bright yellow-green and olive-green points show the probabilities of reemergence in ≤ 40 years for DENV strains in *w*AlbB and *w*MelM, respectively, with the size of the point reflecting the number of strains with each probability of reemergence.

We next modelled the probability of DENV reemergence within a 40 year period for *w*AlbB and *w*MelM (**Fig. 6d**), given the specific relative dissemination levels observed in the experimental data. We found the greatest probabilities of reemergence for strains of DENV-3 under *w*AlbB. We estimated a high probability of reemergence for at least one DENV-3 strain in settings with a baseline R_0_ as low as 1.5, while at R_0_>3.5 we found 8 of the 11 DENV-3 strains reemerged in all our simulations. We found similar results assuming a *w*MelM intervention, estimating high probabilities of reemergence for half of the modelled strains at R_0_>3.5 and a high probability of reemergence for two strains at an R_0_ of 1.5. Under *w*AlbB, we estimated high probabilities of reemergence for two DENV-1 strains for a baseline R_0_ as low as 2, while at higher R_0_ values we estimated high probabilities of reemergence for around a third of modelled DENV-1 strains. When considering *w*MelM, we only found high probabilities of reemergence of around two DENV-1 strains at high baseline R_0_s. Overall, we found little evidence for reemergence of DENV-2 strains under either *w*AlbB or *w*MelM. We estimated a high probability of reemergence of only one DENV-2 strain under *w*AlbB assuming baseline R_0_s of >4 and no reemergence of DENV-2 strains under *w*MelM at any assumed baseline R_0_. Similarly, we found no evidence of reemergence of DENV-4 under *w*AlbB or *w*MelM under any assumed baseline R_0_.

Finally, we used a multi-strain version of the model (assuming five circulating strains per serotype) to estimate the impact of differential *Wolbachia* inhibition on strain diversity (**Fig. 6b**). We estimated the average dissemination of all strains, weighted by their prevalence and used a threshold of 0.1 per 100,000 population to define whether a strain was circulating. We found evidence for selection for DENV strains with higher relative dissemination over time in both *w*AlbB and *w*MelM scenarios. In the *w*AlbB scenario, there was initially a median average relative dissemination of 0.27 (95% CrI: 0.13-0.40), while by the end of the 75 year simulation period a median of three strains remained in circulation with an average relative dissemination of 0.57 (95% CrI: 0.57-0.58). Similarly, for *w*MelM the initial median average relative dissemination of 0.13 (95% CrI: 0.025-0.32) increased to 0.77 (95% CrI: 0.77-0.77) with a median of one remaining circulating strain.

## Discussion

The release of *Wolbachia*-transinfected *Ae. aegypti* mosquitoes has been shown to effectively inhibit DENV transmission and reduce the public health burden caused by dengue^8,9,31^. Here, we determined the impact of DENV genetic diversity on inhibition by two commonly used *Wolbachia* strains (*w*AlbB and *w*MelM), using a panel of 60 genetically diverse DENV isolates. We combined our experimental findings with transmission dynamic models to elucidate the robustness of *Wolbachia* control of the genetically diverse DENV. Our findings showed that the genetic diversity of DENV impacts the outcome of *Wolbachia* inhibition in mosquitoes, with heterogeneity in the level of inhibition between the two *Wolbachia* strains and the DENV serotypes. In particular, we found an overall stronger inhibition by the *w*MelM strain as compared to *w*AlbB, as well as DENV serotype-specific effects with the least inhibition of many DENV-3 isolates by both *Wolbachia* strains. Furthermore, our models highlight the potential for DENV strain selection and demonstrate that the probability of DENV reemergence is dependent on the assumed transmission intensity, deployed *Wolbachia* strain, and circulating DENV serotype.

Previous studies on vector competence of wildtype *Ae. aegypti* mosquitoes for DENV have typically focused on a relatively small number of viruses^32^. Our approach of including a large panel of 60 genetically characterized and diverse DENV isolates is a significant advancement in the field and allows for direct comparisons within and between DENV serotypes, in the presence and absence of *Wolbachia*. While keeping the genetic background of the mosquito constant, we found that exposing wildtype *Ae. aegypti* to our panel of DENV isolates exhibited a high degree of variation in infection and dissemination rates ranging from 0% to 100%, even in absence of *Wolbachia* (**Fig. 2**). This shows that genetically diverse DENV isolates can have significantly different outcomes on vector competence in *Ae. aegypti*. Indeed, previous studies have arrived at similar findings, but only included a few selected DENV isolates per serotype^25,33^. For the first time, we have performed a vector competence study representing the majority of genotypes within each of the four DENV serotypes, assessed side-by-side in *Ae. aegypti* mosquitoes. Moreover, for several genotypes, including genotypes 1I, 1V, 2II, 3III and 4I, we included at least 5 DENV isolates per genotype, which revealed variation in DENV infection and dissemination not only at the serotype level, but also at the genotype level. Our findings on the susceptibility of wildtype *Ae. aegypti* for genetically diverse DENV isolates provide important new insights in the variability within DENV serotypes and genotypes at an unprecedented resolution.

By including two *Wolbachia* strains (*w*AlbB and *w*MelM) in our experimental studies, we were able to assess *Wolbachia*-strain specific impacts on inhibition of all four DENV serotypes and their respective genotypes. Overall, we found stronger inhibition of DENV by *w*MelM than *w*AlbB, but the degree of inhibition depended on specific interactions between *Wolbachia* strains and DENV serotypes. The stronger inhibition by *w*MelM was particularly notable for many DENV-2 isolates, where dissemination was completely inhibited for most of the genotypes by *w*MelM, whereas we observed intermediate inhibition of all genotypes by *w*AlbB (**Fig. 2**, **Fig. 3b**). In contrast, the level of inhibition of the other three serotypes was more similar between both *Wolbachia* strains (**Fig. 3b**, **Supplementary Fig. 2**). Relatively few studies have compared the ability of the *w*AlbB and *w*Mel strains to inhibit a DENV under the same experimental design^19,20,26,34^. Most of these studies determined inhibition of DENV-2 isolates and their findings align with our DENV-2 results, but they were not able to reveal the intricate interactions with all four DENV serotypes as we observed in our study. Despite the overall stronger inhibition by *w*MelM, *w*AlbB may remain important to the field due to its high heat tolerance^35,36^. High temperatures differentially impact *Wolbachia* strains, particularly *w*Mel, by lowering *Wolbachia* densities and virus inhibition^36–38^. Importantly, the *w*MelM variant of *w*Mel that we used in our study has increased heat tolerance compared to the other *w*Mel variant being released for DENV control and could provide a solution under warmer climates^19,39^. Our findings highlight the need for future studies aimed at uncovering the even more complex interactions between DENV genotype, *Wolbachia* genotype, *Ae. aegypti* genotype, and environmental factors, such as temperature (i.e., G x G x G x E interactions).

Our work also demonstrates the variation in *Wolbachia* inhibition between and within DENV serotypes, with several DENV-3 isolates less likely to be inhibited by both *w*AlbB and *w*MelM (**Fig. 2b**, **Fig. 3b**). Given the large number of DENV isolates represented in our study, we show that a high level of heterogeneity of *Wolbachia* inhibition exists within DENV-1, 3, 4, and to a certain degree in DENV-2. This highlights that selection of few “representative” DENV isolates per serotype in experimental studies may not accurately represent the heterogeneity in inhibition that may exist within each serotype. For example, if we had chosen DH102 (low relative dissemination in both *w*AlbB and wMelM) to represent DENV-3 or genotype 3III, then we would conclude strong inhibition by *Wolbachia*, whereas selecting DH168 (high relative dissemination in both *w*AlbB and *w*MelM) would have resulted in the opposite. While we found clear differences in inhibition expressed as reductions in infection and dissemination rates, we did not observe strong reductions in DENV replication in fully disseminated mosquitoes (**Fig. 5a**). This is in contrast to prior studies that reported strong reductions in DENV replication in *Wolbachia*-transinfected mosquitoes^40^. Overall, our comprehensive approach revealed how variable *Wolbachia* inhibition can be between DENV serotypes, genotypes, and isolates.

Mathematical models allowed us to understand the implications of these findings on long-term population effects of release programs, demonstrating higher risk of reemergence for DENV-3 strains across a range of plausible transmission intensities under both *w*MelM and *w*AlbB, as well as reemergence of DENV-1 strains under *w*AlbB. Contrastingly, we found little or no evidence for reemergence of nearly all DENV-2 and DENV-4 strains considered. This has potential epidemiological significance depending on the strains that are circulating or introduced in real-world settings. Additionally, we demonstrated the potential for DENV strain selection following *Wolbachia* interventions in both *w*AlbB and *w*MelM scenarios, with selection for a median of three strains under *w*AlbB and one strain under *w*MelM with high average relative dissemination by the end of the simulation period. Taken together, these results demonstrate that differential inhibition of *Wolbachia* on DENV strains (combined with an accumulation of susceptible individuals following a *Wolbachia* intervention reducing DENV transmission) could allow for reemergence of DENV and selection of strains that are least inhibited by *Wolbachia.* This would reduce the potential impact of the intervention in the medium-to-long term and has important implications for the planning of release programs.

While our study provides important new insights in the impact of DENV genetic diversity on inhibition by *Wolbachia*, it does have some limitations. First, artificial membrane feeding systems, although ideal for our controlled experiment, are not a perfect representative of mosquito blood-feeding in nature^41,42^. However, for the purpose of our study, it was necessary to standardize the conditions in which our three *Ae. aegypti* colonies were exposed to each DENV strain. Second, we used DENV virus directly harvested from cell culture (i.e. without freezing for storage) to prepare infectious blood meals, following previous work that shows that freezing DENV stocks negatively impacts mosquito infection rates^29^. While this was critical to achieve sufficient infection rates, it resulted in variability in DENV titers between strains. We addressed this limitation by exposing WT, *w*AlbB-, and *w*MelM-transinfected in parallel to each DENV stock, and by determining the relative dissemination rate of each *Wolbachia* colony compared to WT control to account for variability in infection in WT mosquitoes. Third, we used dissemination as a proxy for transmission, instead of detecting DENV presence in saliva, as a previous study demonstrated that it is a more accurate estimate of transmission to a host than forced salivation assays^24^. Fourth, our mathematical modelling framework relied on several assumptions. In particular, we calibrated our model using historical and projected demographic data for Brazil, a dengue-endemic country. As such, the results may not be representative for settings with different levels of endemicity. We also assume complete *Wolbachia* introgression with a country-wide roll-out and do not consider the additional impacts of patchy or imperfect *Wolbachia* implementation on disease burden^44^. Additionally, we use data from the large number of DENV isolates represented in this experimental work to parameterise our models, which may not be representative of the strain diversity observed in endemic settings. However, this also allowed us to characterise a broad range of potential epidemic dynamics that could be observed under *Wolbachia,* which would not have been possible with a smaller sample of DENV isolates.

Despite these limitations, our study makes a strong case for considering DENV genetic diversity when implementing *Wolbachia*-based control interventions. *Wolbachia* release programs already take several ecological factors into account before release, such as the inhibition strength, host fitness costs, thermal stability of specific *Wolbachia* strains, and backcrossing *Wolbachia* into locally collected mosquito populations to match the environment of release^45,46^. Despite these considerations, there is variation in the level of dengue case reduction that is achieved in different field trials, particularly in complex urban settings^46,48^. Based on our laboratory findings, we recommend broadly implementing DENV genomic surveillance in *Wolbachia* field trials as an important next step to further improve our understanding of the role of virus genetic diversity in *Wolbachia*-based control of dengue.

## Methods

### DENV isolates

To create a comprehensive panel of genetically diverse DENV isolates, we selected a subset of viruses from collections provided by the Yale Arbovirus Research Unit and World Reference Center for Emerging Viruses and Arboviruses. From this collection, 9-20 of the most diverse isolates were selected in an attempt to represent each of the genotypes within each serotype. All major genotypes, except 1VI, 3V, and 4III, for which no virus stocks were available, were included in the final panel; for genotypes with larger clades, multiple representatives with the highest genetic divergence were chosen. Additional considerations included the selection of isolates with the lowest number of prior passages in cell culture. This resulted in a final selection of 20 DENV-1 (genotypes I-V, and VII), 20 DENV-2 (genotypes I-VI), 11 DENV-3 (genotypes I-III), and 9 DENV-4 (genotypes I, II, and IV; **Fig. 1a-b**; **Supplementary Table 1**). All lyophilized virus stocks were resuspended and passaged three times on *Ae. albopictus* C6/36 cells to expand stocks and achieve sufficient virus titers. After five days of inoculation, third passage viruses were harvested, centrifuged for 10 min at 3200 x g at 4°C to remove cell debris, and directly used in mosquito infection experiments. All remaining virus stock aliquots were stored at -80°C. C6/36 cells were maintained by splitting every 3-4 days at a 1:15 dilution using MEM media with 10% fetal bovine serum, 1% antibiotic-antimycotic, and 1% non-essential amino acid.

### Virus metagenomic sequencing

Given that virus stocks originated from several different sources and had varying passage history, we used metagenomics for whole genome sequencing to verify the identity and purity of each stock after two initial passages. This confirmed the identity of the 60 DENV isolates in our panel (**Fig. 1b**), and revealed that two DENV stocks (DH069 and DH169) had insect-specific viruses present, for which we were not able to select an alternate genetically related stock (**Supplementary Table 1**; **Supplementary Data File 1**). Viral RNA was extracted using the MagMAX Viral/Pathogen Nucleic Acid Isolation Kit on the KingFisher Flex purification system. Sequencing libraries were prepared using an untargeted metagenomic approach adapted from Matranga et al. 2014^49^. For each sample, 20 μL of RNA was DNase-treated using 3.5 μL DNase I reaction buffer (10X), 0.5 μL DNase I, 0.5 μL RNase Inhibitor, 0.5 μL linear acrylamide, and 10 μL nuclease-free water. The reaction was incubated at 37°C for 30 minutes. RNA was purified using a ratio of 1.8:1x volume Mag-Bind Total Pure NGS beads:sample. First strand cDNA was synthesized using 10 μL of DNase-treated RNA, 4 μL of SuperScript IV VILO master mix, and 6 μL of nuclease-free water. The reaction was incubated at 25°C for 10 minutes, 50°C for 10 minutes, and 85°C for 5 minutes. Second strand cDNA was synthesized by combining 20 μL of first strand cDNA, 8 μL of 10X NEB Second Strand Reaction Buffer, 3 μL dNTP mix (10 mM), 1 μL DNA Ligase (10 U/μL), 4 μL DNA Polymerase I (10 U/μL), 1 μL RNase H (2 U/μL), and 43 μL of nuclease-free water. The reaction was incubated at 16°C for 2 hours, inactivated by adding 5 μL of 0.5M EDTA, followed by purification using the same 1.8:1 ratio. The Illumina Nextera XT DNA Library Preparation Kit was used to prepare DNA libraries following manufacturer’s protocols with half the recommended reaction volumes, except for the cDNA input and Amplicon Target Mix where we used 4 μL and 1 μL, respectively. Final libraries were quantified using the Qubit 1X dsDNA HS Assay Kit following the manufacturer’s protocol. Libraries were normalized and subsequently pooled for sequencing on the Illumina NovaSeq platform with paired-end reads of 150 base pairs in length and targeting 2 million reads per sample at the Yale Center for Genome Analysis. Two negative (nuclease-free water) controls were included in the library preparation process.

Consensus genomes were generated using iVar with a minimum frequency threshold of 75% and minimum depth of 10 reads. Consensus genomes were submitted to NCBI GenBank and raw sequencing data to Sequence Read Archive (SRA) under BioProject number PRJNA1308108. The CZID metagenomics pipeline for Illumina and Nanopore sequencing platforms was used to verify identity and purity (e.g. detect the presence of other viruses, such as insect-specific viruses) of all DENV stocks (**Supplementary Data File 1**)^50^. Water controls were used as background models in the analysis. Sequencing reads were filtered for non-phage viruses and using thresholds of NT rPM (normalized reads per million) ≥ 10, NT L (alignment length in bp) ≥ 50, and NT Z-score ≥ 2 to ensure detection of sequences significantly different from background noise.

### Mosquito rearing

Three *Aedes aegypti* colonies (wildtype control (WT; no *Wolbachia*)), or transinfected with *Wolbachia w*AlbB or *w*MelM strain) with similar genetic backgrounds were used in this study. WT mosquitoes were originally collected from wild populations in Cairns, Australia. *w*AlbB mosquitoes are WT mosquitoes stably transinfected with *Wolbachia* from *Ae. albopictus* mosquitoes and *w*MelM mosquitoes are WT mosquitoes stably transinfected with *Wolbachia* from *Drosophila melanogaster*^51^. Both *Wolbachia* strains have been characterized previously and released in field trials^52^. Each mosquito colony was maintained in separate environmental chambers at 26°C, 60-70% relative humidity, and 12:12 light-dark cycle^26^. Eggs were added to trays with 500 mL of Picotap water and 2 drops of Liquifry No.1 fish food. After hatching, approximately 200 first-instar larvae were transferred to a tray with 1 L water and fed with Tetramin baby fish food. Pupae were collected in small cups and transferred to BugDorm-1 cages. Adult mosquitoes had unlimited access to a 10% sucrose solution and were blood-fed with defibrinated sheep blood using a Hemotek membrane feeder for egg production. For mosquito infections, 1-2 week old adult female mosquitoes were counted and transferred to cups and kept on a 10% sucrose solution. One day prior to infectious blood meals, 10% sucrose was replaced with water to stimulate blood feeding the next day.

### Mosquito infections

To determine inhibition of our DENV stocks by *Wolbachia*, we exposed our WT, *w*AlbB, and *w*MelM colonies in parallel to infectious blood meals using the Hemotek artificial membrane feeding system. Directly after DENV stocks were harvested on day five after inoculation, infectious blood meals were prepared by mixing DENV 1:1 with defibrinated sheep blood. Each DENV-blood mixture was loaded into three Hemotek reservoirs and sealed with a Parafilm membrane to allow parallel feeding of the three *Ae. aegypti* colonies from the same virus stock. Sixty mosquitoes were allowed to feed for 1-1.5 hours, immobilized on ice, and engorged females were collected and placed back in cups for 10 days incubation at 26°C. Given the large number of 180 unique groups (3 mosquito colonies x 60 DENV isolates), the feeding experiment was performed in three different batches, with 10 viruses tested in each of the three mosquito colonies on two separate days per batch (30 combinations per day; 60 combinations total per batch). All twenty DENV-1 viruses were tested in batch 1, all twenty DENV-2 viruses were tested in batch 2, and all eleven DENV-3 and nine DENV-4 viruses were tested in batch 3. To account for possible batch effects, each experimental day included a DENV-2 virus (DH069) as a control that was fed to WT mosquitoes. This batch control showed that there was variation in infection and dissemination between batches, which correlated with variation in the input stock virus titer used to prepare the infectious blood meal (**Supplementary Fig. 4**).

All mosquitoes were dissected 10 days after incubation, with bodies and legs collected in separate tubes. Each tube contained a 3.2 mm stainless steel bead and 200 µL of mosquito diluent (1X phosphate buffered saline, 20% fetal bovine serum, 50 µg/mL Penicillin-Streptomycin, 50 µg/mL of Gentamicin, and 2.5 µg/mL of Amphotericin B). All samples were stored at -80°C until further processing. Mosquito bodies and legs were homogenized using the Bullet Blender Storm Pro Bead Homogenizer for 2 minutes at speed 8, and centrifuged for 1 minute at 14,500 rpm. Infection (presence of DENV in mosquito bodies) and dissemination (presence of DENV in mosquito legs) rates were determined using RT-qPCR.

### DENV RT-qPCR

Total nucleic acid was extracted from all mosquito bodies and a subset of legs using the NEB Monarch Mag Viral DNA/RNA Extraction Kit on the KingFisher Flex purification system. Mosquito homogenates (75 µL) were eluted into 75 µL elution buffer for each reaction. A direct pairwise comparison was made between RT-qPCR cycle threshold (Ct) values of mosquito homogenates with and without RNA extraction. This revealed that RNA extraction of mosquito legs could be omitted without loss of sensitivity (**Supplementary Fig. 5b**), while some variation was observed when testing mosquito bodies without RNA extraction (**Supplementary Fig. 5a**), which still warrants further optimization.

Mosquito bodies were tested for the presence of DENV using the NEB Luna Universal Probe One-Step RT-qPCR kit, with previously designed gBlock positive controls and a primer and probe set that targets all four DENV serotypes (**Supplementary Table 2**)^53^. The primers and probes were aligned to 59 genomes with sufficient coverage of the 3’ UTR site to verify conservation of the binding regions. The RT-qPCR protocol was run on a Bio-Rad CFX Opus real-time PCR system using the following conditions: 55°C for 10 minutes, 95°C for 1 minute, followed by 40 cycles of 95°C for 10 seconds, 60°C for 30 seconds, and a plate read. Samples with a Ct value lower than 38 were considered DENV positive. Mosquito legs of bodies that tested positive were directly tested by RT-qPCR without prior RNA extraction. Each qPCR run included a 10-fold serial dilution of gBlock controls (1 to 10^6^ genome equivalents (GE)/µL) along with a no-template negative control. The resulting standard curve generated from Ct value and gBlock copy number was used to convert Ct values to estimate genome equivalents in positive mosquito samples.

### Focus forming assay

Focus forming assays (FFA) were used to determine infectious titers of DENV stocks used in mosquito infection experiments, following the protocol by Vasilakis et al. (2009)^54^. Each virus stock was serially diluted from 1e-2 to 1e-5. A total of 100 µL of each dilution was inoculated on 80% confluent Vero cells in 24-well plates. Plates were rocked every 20 mins for 2 hours total before being overlaid with 1 mL of methylcellulose. After 5 days of incubation at 37°C, cells were fixed with cold 90% methanol, incubated for 50 minutes at room temperature and methanol was then discarded and air dried for 50 minutes. Plates were incubated with blotto (3.5% milk in 1X PBS) to block non-specific binding. Primary and secondary antibodies were diluted in blotto at 1:1000. Fixed plates were incubated with 200 µL antibodies for 1.5 hours, at 37°C. Mouse anti-DENV type 1,2,3,4 monoclonal primary antibodies (Creative-Diagnostics, #DCABH-10243) were used for DENV-1, DENV-2, and DENV-3, but did not work for DENV-4 viruses. Thus, monoclonal anti-dengue virus type 4 envelope protein antibodies clone E62 (BEI Resources, NR-15551) were used for DENV-4. Primary antibodies were discarded, then plates were washed with 1 mL 3.5% blotto. Secondary antibody staining (using SeraCare #5220-0339 antibodies for all samples) was then performed by incubating for 1 hour with 200 µL human serum adsorbed and peroxidase-labeled anti-Mouse IgG (gamma) antibodies. After discarding secondary antibodies, plates were washed with 1X PBS and incubated with 100 µL Trueblue peroxidase substrate for 12 minutes while covering to protect from light. Wells with manageable number of foci (approximately 20-90) were counted in replicate, averaged, then multiplied by dilution factor and divided by the volume of inoculum (100 µL) to determine the focus forming unit per mL (FFU/mL). RT-qPCR and FFA results were compared for all DENV stocks used in mosquito experiments (**Supplementary Fig. 6**). Three DENV-4 viruses were below the FFA detection limit.

### *Wolbachia* relative density

After nucleic acid extraction, a subset of 178 mosquito bodies were screened to determine the relative density of *Wolbachia* by normalizing *Wolbachia* copy numbers to *Aedes* ribosomal protein S6 housekeeping gene copy numbers. The NEB Luna Universal qPCR Master Mix kit was used to determine the DNA copies for both *Wolbachia* and *Aedes*, using previously published primers (**Supplementary Table 2**)^55^. The RT-qPCR protocol was run on a Bio-Rad CFX Opus real-time PCR system using the following conditions: 95°C for 3 minutes, followed by 40 cycles of 95°C for 10 seconds, 60°C for 30 seconds; a melting curve from 65-95°C with a rate of 0.5°C every 5 seconds was used to determine the melting point of any amplified DNA. For the *Wolbachia* relative density (*w*M*w*A) assay, WT mosquitoes had no melt peak, *w*AlbB mosquitoes had a melt peak of 78-78.5°C, and *w*MelM mosquitoes had a melt peak of 80.5°C. For the *Aedes* assay, all samples (WT, *w*AlbB, and *w*MelM) had melting peaks between 81.5-82.5°C. The relative density of *Wolbachia* was calculated using the formula 2^(*Aedes* Ct - *w*M*w*A Ct)^56^.

### Data analysis

Phylogenetic trees for each of the four DENV serotypes were constructed using the Nextstrain tool Augur^57,58^. Initial visualization was done on auspice after which Newick tree files were exported and a final tree visualization highlighting selected viruses was generated using Baltic^59^.

Infection rate was calculated as the percentage of virus-positive mosquito bodies (Ct < 38) out of the total number of exposed mosquitoes, and dissemination rate was calculated as the percentage of virus-positive legs (Ct < 38) out of the total number of exposed mosquitoes. Dissemination rates were used as a proxy for virus transmission, as previous studies found that testing mosquito legs was more reliable than saliva collection^24^. Relative infection for each virus was calculated as the ratio of infection rate in *Wolbachia* mosquitoes to that in wildtype mosquitoes. Similarly, the relative dissemination for a virus was determined as the ratio of dissemination rate in *Wolbachia* mosquitoes to that of wildtype mosquitoes. Statistical differences in relative infection and dissemination between *Wolbachia* strains and DENV serotypes were determined using the Wilcoxon test for pairwise comparisons. To account for multiple comparisons, we applied the Benjamini-Hochberg correction to control the false discovery rate. The Pearson correlation test was used to assess relationships. We used the R statistical software (v4.0.2) for all statistical analyses. Code used for the statistical analyses are publicly available on https://github.com/vogels-lab/2025_paper_denv-wolbachia-genetic-diversity.

### Transmission dynamic model

To evaluate the potential impacts of the observed diversity in *Wolbachia* inhibition, we used a multi-strain stochastic compartmental model to simulate dengue epidemic dynamics in *Wolbachia* intervention scenarios. Within our modelling framework, individuals can be infected four times (once with each DENV serotype). We assume homotypic immunity is lifelong and heterotypic immunity lasts one year^60^. The model is stratified into single-year age groups (up to 80, then 80+) and keeps track of an individual’s immune history. We track which serotypes individuals have been infected with but not the order of past infections. We use two versions of the model in our simulation experiments: one assuming a single circulating strain per-serotype and one assuming five circulating strains per-serotype (and so including strain competition). We do not explicitly model mosquito dynamics, instead modelling seasonality in transmission using a cosine function in our force of infection term. Our model follows a cohort ageing process, where individuals move up an age class once per year. Full details on model equations and parameters can be found in the **Supplementary Materials**.

We calibrated the model using historical demographic data for Brazil from the UN’s World Population Prospects 2024, available from 1950 to 2023^61^. We calibrated the model for 100 years, assuming static demography prior to 1974. We repeated model calibration 50 times, generating 50 sets of initial conditions for onward simulation. We ran model calibration separately for two versions of the model: a four serotype model and a 20 strain model (with 5 strains per serotype).

We then simulated potential epidemic dynamics under *Wolbachia* interventions over a 75 year horizon from 2024 to 2100. We conducted two sets of simulations. In the first we used a serotype-level model to estimate the probability of serotype reemergence under partial *Wolbachia* inhibition. We assumed complete *Wolbachia* introgression from the first day of the simulation as well as total inhibition of three serotypes by *Wolbachia* and partial inhibition of the fourth. We parameterised inhibition by *Wolbachia* as multiplier on the transmission coefficient (β), and considered different levels of relative dissemination in *Wolbachia*-infected mosquitoes compared to wildtype. We ran simulations considering R_0_s ranging from 1 to 5 in 0.5 increments and relative dissemination ranging from 0 to 1 in 0.05 increments. For each of these 189 parameter combinations we ran 50 stochastic simulations, each using a different set of initial values. We then repeated this, varying the serotype experiencing partial inhibition, resulting in 200 stochastic simulations per parameter set. We used a threshold of 1 infection per 100,000 people to define extinction and re-emergence of DENV following the implementation of a *Wolbachia* intervention and estimated the time until re-emergence for each simulation.

We then ran simulations considering levels of inhibition corresponding to the relative dissemination seen for the DENV virus strains in *w*AlbB and *w*MelM mosquitoes in **Fig. 3**. We ran 50 stochastic simulations for each parameter set. From these, we estimated the probability of reemergence in 40 years for each DENV strain as the number of simulations in which reemergence occurred within 40 years divided by the total number of simulations.

Finally, we used a multi-strain version of the model to simulate potential DENV ecological dynamics under *Wolbachia* interventions. Here, we drew five strains per serotype and their corresponding relative dissemination under *w*AlbB and *w*MelM (as in **Fig. 3**). We repeated this 20 times with replacement and ran 50 stochastic simulations per strain draw (resulting in 1000 simulations in total). We then simulated strain dynamics over time and estimated the average relative dissemination of circulating strains, weighted by their prevalence. We used a threshold of 0.1 infections per 100,000 to define whether a strain was circulating.

### Data availability

All data are included in this manuscript, the supplementary information, and on BioProject PRJNA1308108. Source data are provided with this paper.

### Code availability

All code and data used for the statistical analyses and transmission modelling analysis can be found at https://github.com/vogels-lab/2025_paper_denv-wolbachia-genetic-diversity and https://github.com/EmilieFinch/denv-wolbachia.

## Acknowledgements

We would like to thank S. Weaver, J. Plante, and K. Plante from the University of Texas Medical Branch for sharing dengue virus stocks from the World Reference Center for Emerging Viruses and Arboviruses, and D. Brackney, A. Bransfield, and P. Armstrong from the Connecticut Agricultural Experiment Station for sharing dengue virus stocks from the Yale Arbovirus Research Unit Collection. We thank Prof. Ary Hoffmann for providing resources critical for mosquito colony production. This publication was made possible by CTSA Grant Number UL1 TR001863 from the National Center for Advancing Translational Science (NCATS), a component of the National Institutes of Health (NIH) awarded to C.B.F.V., the Ambrose Monell Foundation (C.B.F.V.), the NIH T32AI055403 (A.S.), the National Science Foundation Graduate Research Fellowship under Grant No. DGE-2139841 (A.S.), the National Institute of Allergy and Infectious Diseases of the NIH award DP2AI176740 (N.D.G.), an Australian Research Council Discovery Early Career Researcher Award DE230100067 funded by the Australian Government (P.A.R), the European Research Council No. 101170844 (H.S.), the Wellcome Trust PCAG/522 (E.F. and H.S.), and the National Institute of General Medical Sciences of the NIH under Award Number 1S10OD030363-01A1 awarded to the Yale Center for Genome Analysis. The contents of this work are solely the responsibility of the authors and do not necessarily represent the official views of NIH.

## Author contributions

A.S., N.D.G., and C.B.F.V. conceptualized and designed the study. P.A.R. and X.G. created transinfected *Wolbachia* (*w*AlbB and *w*MelM strain) mosquito colonies. A.S., M.I.B., E.B., B.L.N., N.M.F., I.M.O., J.W.B., and C.B.F.V. performed the experiments. A.S., E.F., K.L., V.H., A.T.H., and H.S. analyzed the data. H.S., N.D.G., and C.B.F.V. supervised the study. A.S., E.F., and K.L. wrote the first draft of the manuscript. All authors read, reviewed, and approved the final manuscript.

## Supplementary Data

**Supplementary Data File 1: Outputs from CZ ID for metagenomic sequencing analysis of DENV stocks** File containing raw outputs in reads per million (NT rPM) for metagenomics performed on all 60 DENV isolates. The first tab includes outputs for DENV-1, second tab includes outputs for DENV-2, and third tab includes outputs for DENV-3 and DENV-4. Threshold was set to include sequences with NT rPM ≥ 10, NT L (alignment length in bp) ≥ 50 and NT Z-score ≥ 2.

## Supplementary Methods

### Model structure

To model the potential epidemiological impacts of imperfect *Wolbachia* inhibition we use a stochastic, age-stratified compartmental model structure. Individuals are classified into Susceptible (S), Infected (I), Cross-protected (C), and Recovered (R) compartments. Individuals are initially susceptible to all serotypes (DENV-1-4). After infection with one serotype we assume individuals experience short-lived cross-protection to all serotypes (with a duration of one year). After this, individuals move to the R compartment, where they remain immune to the homologous serotype, but are susceptible to infection by heterologous serotypes. We assume individuals can experience four infections, one per serotype. All compartments are stratified by age group *i* (with 81 age classes from 0 to 80+). I, C and R compartments are also stratified by immune history Ω, keeping track of the set of serotypes an individual has been exposed to (but not the order of infection). Finally, the I compartment is also stratified by the current infecting strain *k*. We use two formulations of the model: the first assumes there is only a single strain per serotype (and so four strains in total). The second assumes there are five strains per serotype (and so 20 in total), in this case an individual can only be infected with one strain per serotype, introducing strain competition.

The model is formulated in discrete time following the equations below. Here δ_*x,y*_ is the Kronecker delta function which is equal to 1 when *x* = *y* and to 0 otherwise. Ω represents the set of serotypes that an individual has been exposed to (but not the order they were experienced in), *a* represents the age class and *k* represents the infecting serotype or strain with total serotypes or strains *K*. *B* represents births, λ_*k*_ represents the force of infection associated with serotype or strain *k*, γ represents the recovery rate and ν represents the rate of loss of cross-protection against heterologous strains.

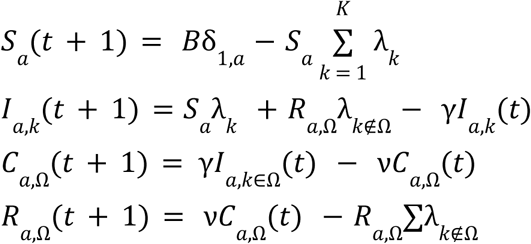

Ageing is modelled as a cohort process, with individuals moving up an age class once per year. At this point, the age class is scaled to match Brazil’s demographic data (i.e., all the compartments for an age class are scaled upwards or downwards to align the total population size of the age class with the demographic data for that year). The total number of births for the year (the population size at evenly throughout the year *a* = 0) are spread out to occur

### Force of infection

For each *R*0 we derived the transmission coefficient β, assuming that each serotype or strain had the same β such that:

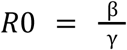

We modelled *Wolbachia* interventions as a reduction on transmission *w* (that is, a multiplier to β) and assumed perfect introgression of *Wolbachia* into the mosquito population on the first day of the simulation (**Supplementary Table 3**). We do not include mosquito dynamics in our modelling framework, instead incorporating seasonality through a cosine function on our force of infection. We also include a λ_*ext*_ parameter to represent external introductions of DENV strains/serotypes.

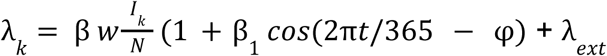

---

**Supplementary Fig. 1.**
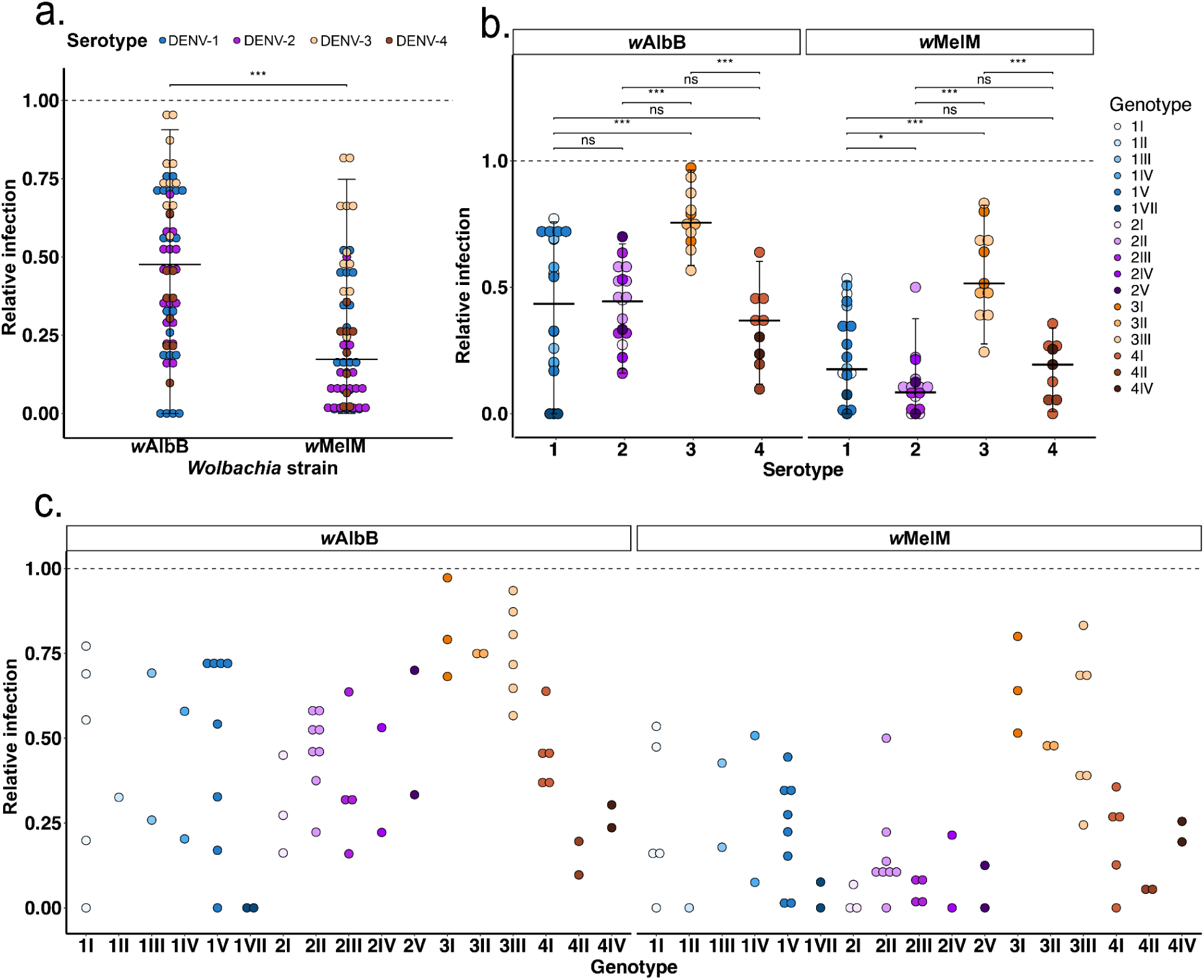
*Wolbachia* strain and dengue virus (DENV) serotype significantly impact the relative infection. Relative infection for each DENV strain was calculated as the ratio of infection rate in *w*MelM or *w*AlbB mosquitoes relative to infection rate in WT mosquitoes. a, The relative infection rate of *w*MelM mosquitoes compared to *w*AlbB for all DENV strains. b, DENV serotype level comparison of relative infection rates for both *w*AlbB and *w*MelM mosquitoes. c, DENV genotype level comparison of relative infection rate for both *w*MelM and *w*AlbB mosquitoes. For figure a-c, each dot represents the relative infection of a DENV strain in either *w*AlbB or *w*MelM mosquitoes relative to WT, with 0 = no infection relative to WT and 1 = equal infection relative to WT. Dots are colored by DENV serotype, with DENV-1 = blue, DENV-2 = purple, DENV-3 = orange, DENV4 = brown. Horizontal bars indicate the median with 95 percent quantile intervals, corresponding to the range between the 2.5th and 97.5th percentiles. The Wilcoxon test was used for statistical comparison with Benjamini-Hochberg correction to account for multiple comparisons. In all panels, *** = p < 0.001, ** = p < 0.01, * = p < 0.05 and ns = p ≥ 0.05. Underlying data for this figure are available in the **Source Data File**.

**Supplementary Fig. 2.**
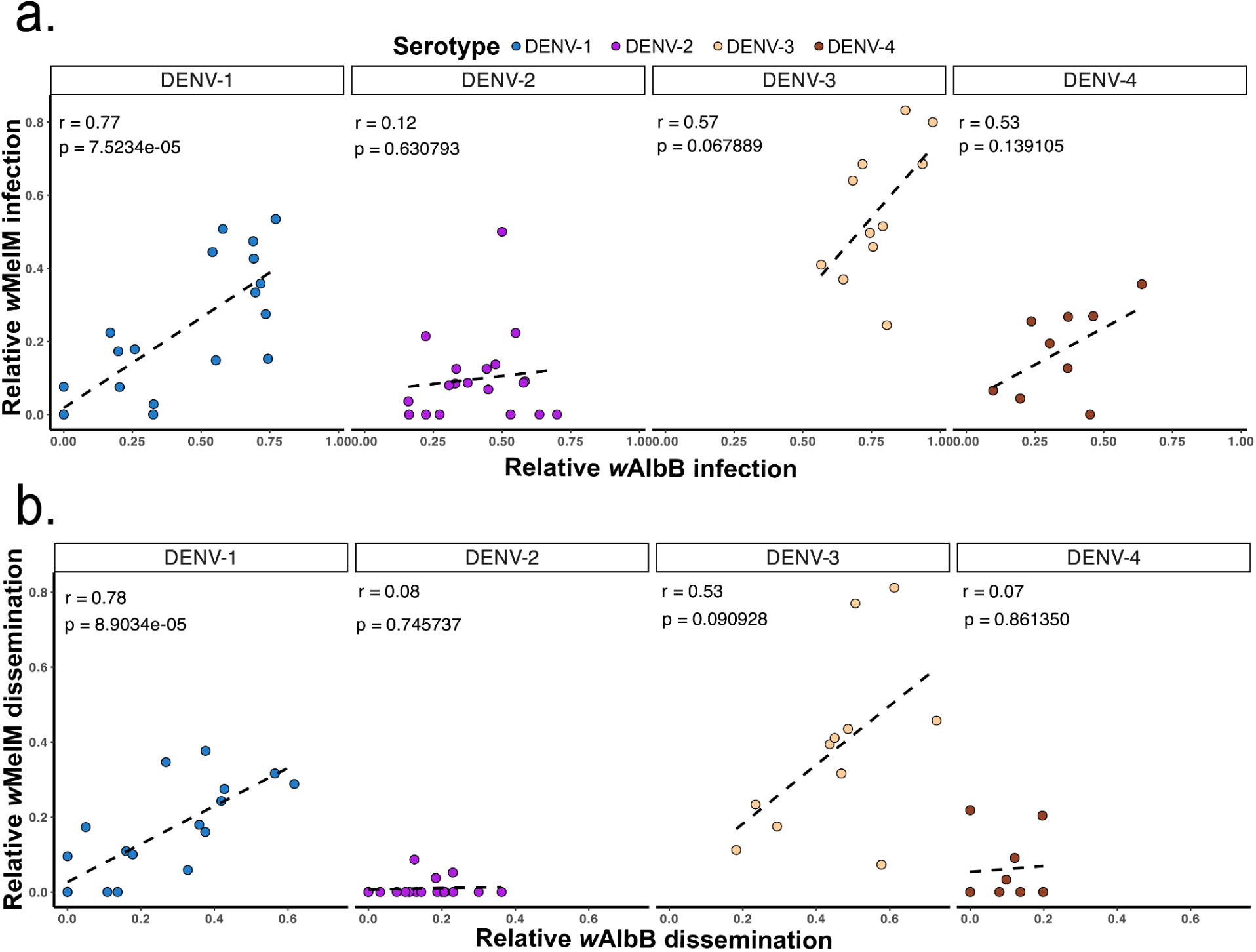
Correlation plots of relative infection and dissemination of *w*AlbB compared to *w*MelM. *Wolbachia* infection and dissemination rates were determined for both *w*AlbB and *w*MelM mosquitoes relative to wildtype (WT). a, Correlation plots comparing relative infection in *w*AlbB compared to relative infection in *w*MelM for each DENV serotype. b, Correlation plots showing relative dissemination in *w*AlbB compared to relative dissemination in *w*MelM for each DENV serotype. a-b, Each dot represents each of the tested DENV isolates, colored by DENV serotype, with DENV-1 = blue, DENV-2 = purple, DENV-3 = orange, DENV4 = brown. The Pearson correlation test was used to assess linear relationships. r = Pearson correlation coefficient and p = p-value. Underlying data for this figure are available in the **Source Data File**.

**Supplementary Fig. 3.**
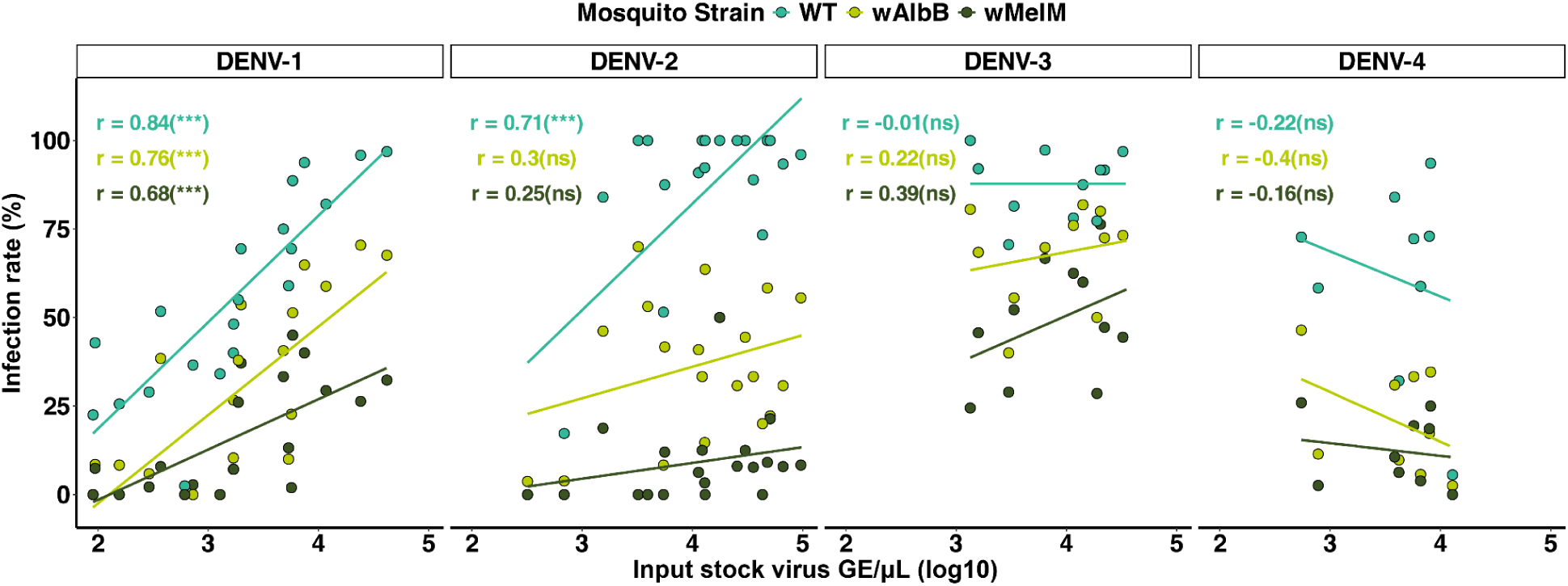
Impact of input stock titer on infection rates across each DENV serotype. Correlation plots of input stock virus genome equivalents with infection rates, colored by mosquito strain, with WT = teal, *w*MelM = olive-green, *w*AlbB = bright yellow-green. Each dot represents each of the tested DENV strains. The Pearson correlation test was used to assess linear relationships. In all panels, r = Pearson correlation coefficient, *** = p < 0.001, ** = p < 0.01, * = p < 0.05 and ns = p ≥ 0.05. Underlying data for this figure are available in the **Source Data File**.

**Supplementary Fig. 4.**
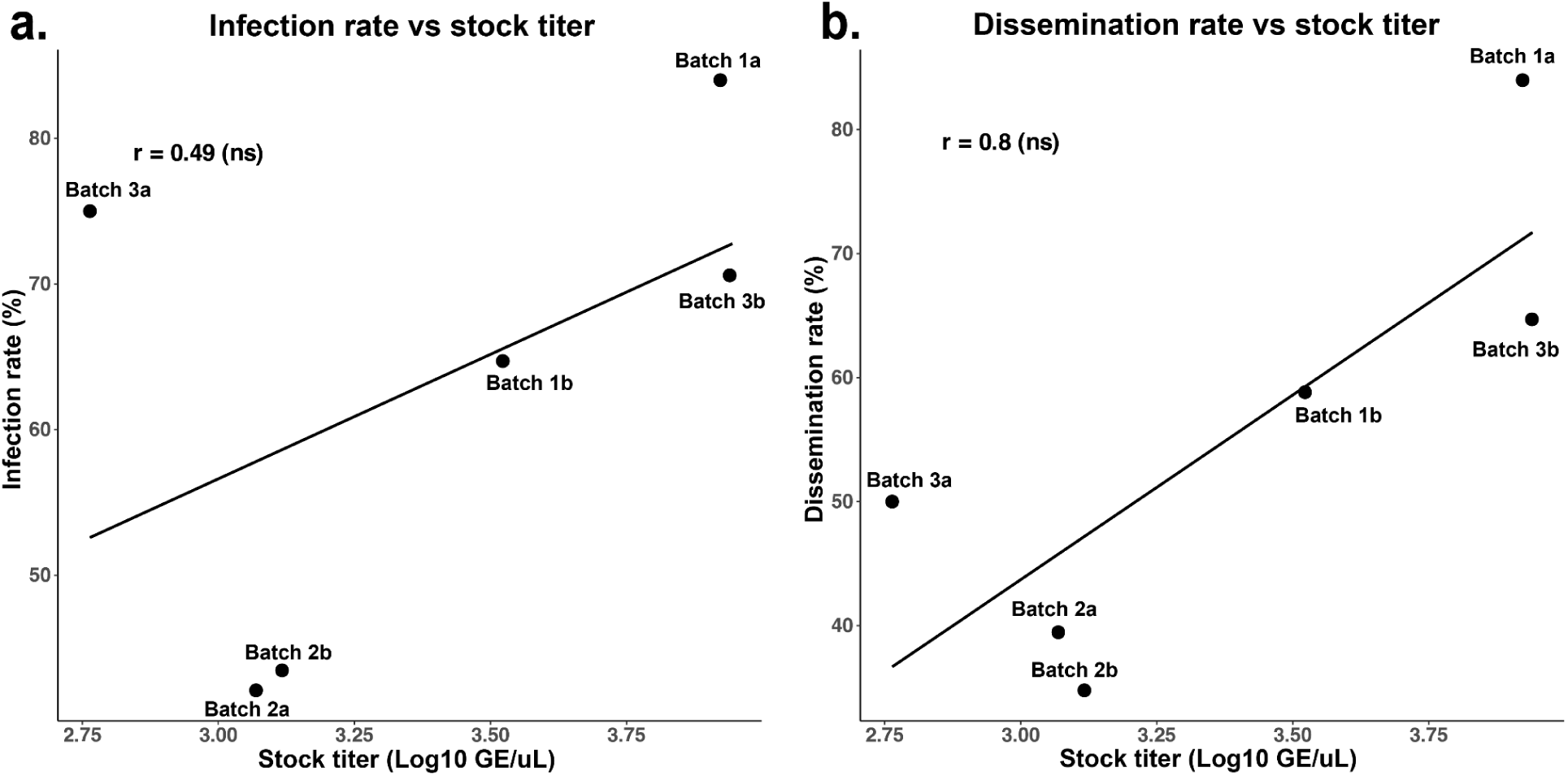
Correlation plots of infection and dissemination rates compared to input stock titer for batch control virus (DH069) exposed to WT mosquitoes. To account for potential batch effects, DH069 virus was included in all experimental batches by exposing WT mosquito colonies to this “control” virus strain. a, Correlation plots comparing infection rate and input stock titer of DH069 for every experimental batch. b, Correlation plots comparing dissemination rate and input stock titer of DH069 for every experimental batch. Each dot represents virus strain DH069 tested in WT mosquitoes across each of the 6 experimental days tested over 3 independent batches. Sample size per dot ranges between 4-38 mosquitoes. The Pearson correlation test was used to assess linear relationships. r = Pearson correlation coefficient and ns = p ≥ 0.05. Underlying data for this figure are available in the Source Data File.

**Supplementary Fig. 5.**
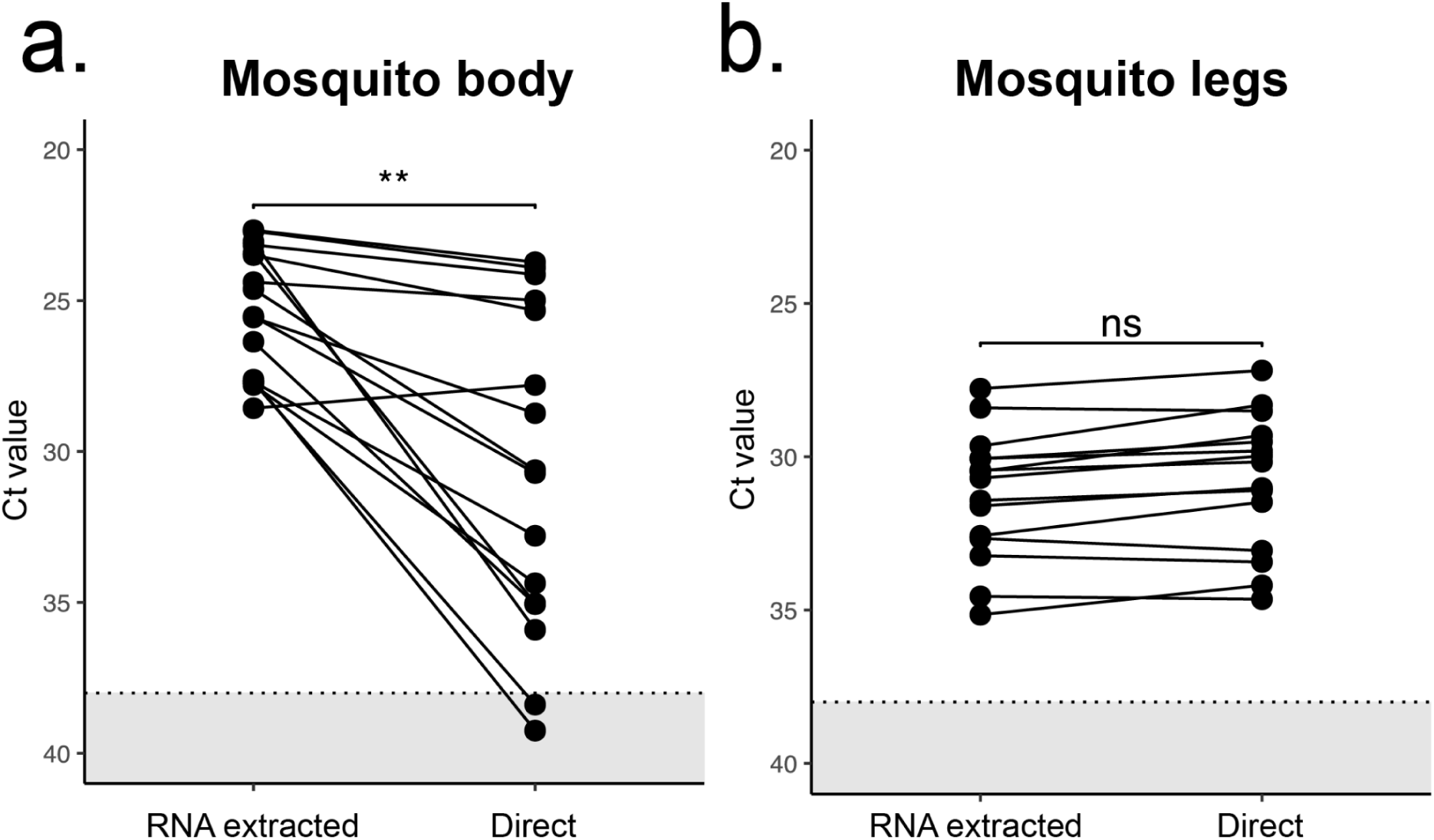
RNA extraction can be omitted when detecting dengue virus (DENV) via RT-qPCR in mosquito legs. Detection of DENV in mosquito bodies and legs spanning a range of Ct values following RNA extraction or directly on homogenates (denoted as “direct”) using RT-qPCR. Dots connected by lines indicate paired RNA extracted and direct samples. a, Comparison between RNA extraction and direct testing of mosquito bodies. b, Comparison between RNA extraction and direct testing of mosquito legs. The Wilcoxon test was used for statistical comparisons. ** = p < 0.01 and ns ≥ 0.05. Samples with a cycle threshold (Ct) value lower than 38 indicated by the dotted line were considered DENV positive. Underlying data for this figure are available in the **Source Data File**.

**Supplementary Fig. 6.**
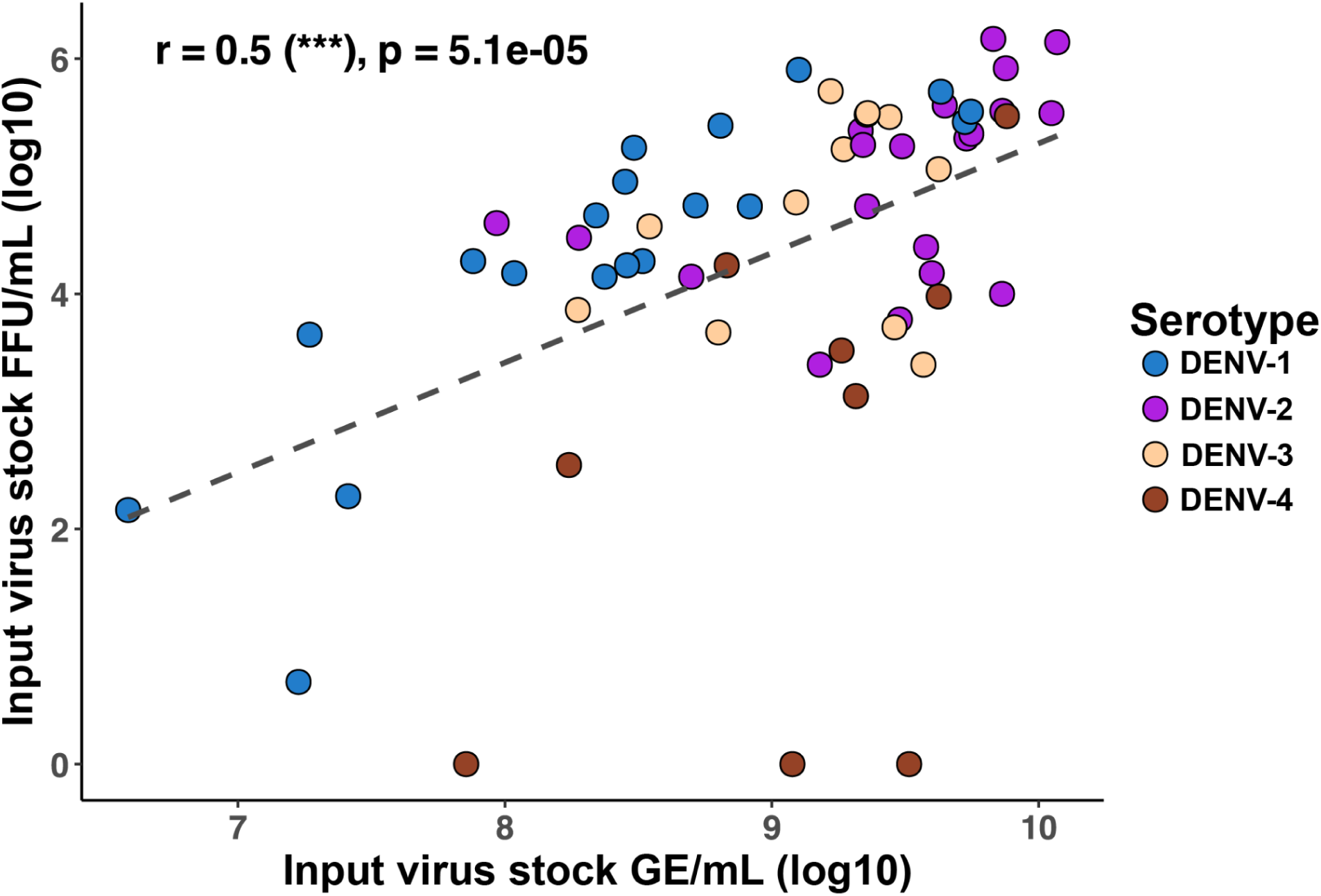
Correlation plot comparing focus forming units to DENV genome equivalents. After the third passage, aliquots of all viruses used in this study were frozen for later quantification. Focus forming assays were used to determine focus forming units and RT-qPCR was used to determine genome equivalents. Each dot represents an individual DENV strain, colored by DENV serotype, with DENV-1 = blue, DENV-2 = purple, DENV-3 = orange, DENV4 = brown. The Pearson correlation test was used to assess linear relationships. r = Pearson correlation coefficient and p = p-value. Underlying data for this figure are available in the **Source Data File**.

**Supplementary Table 1.**
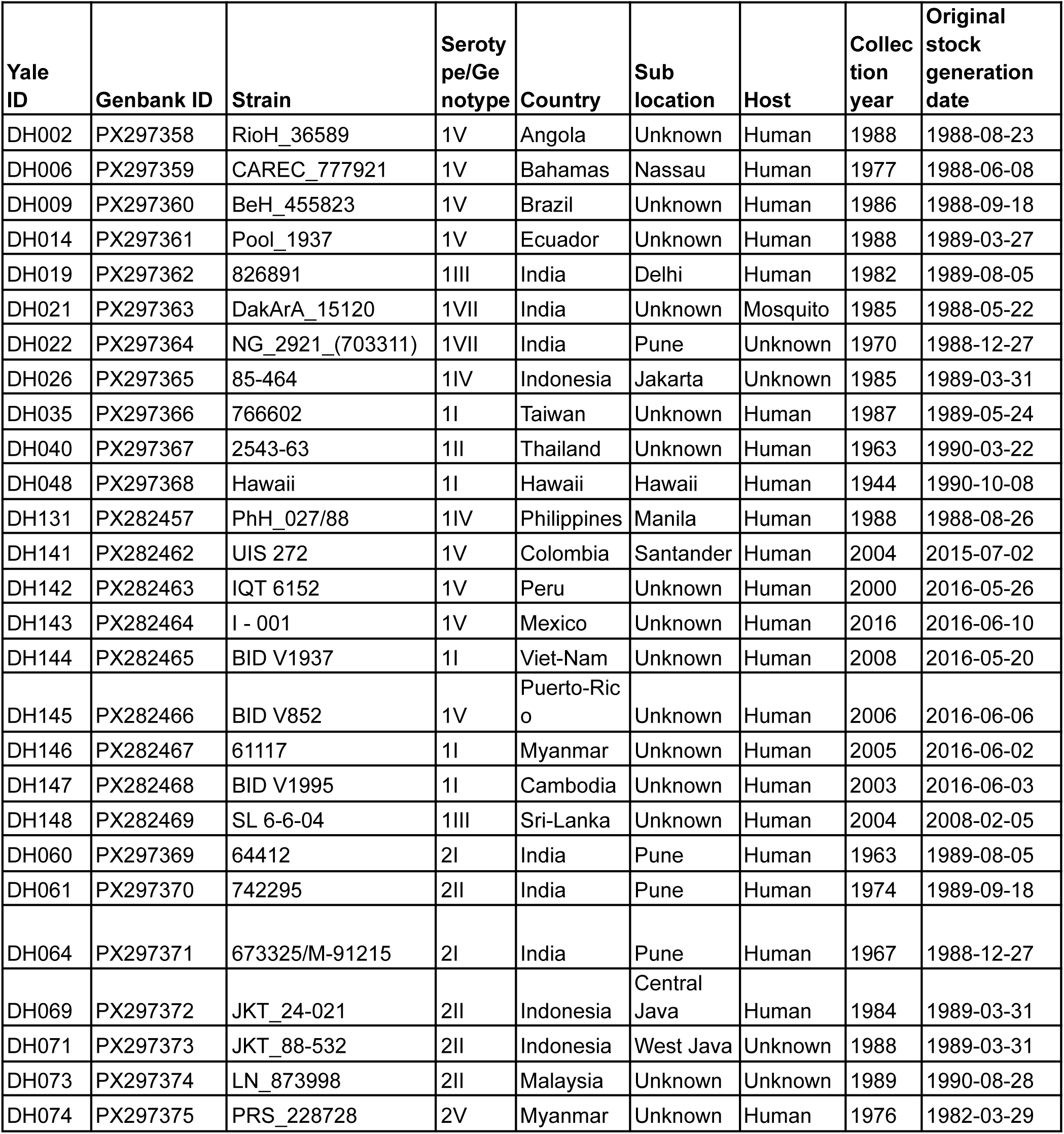
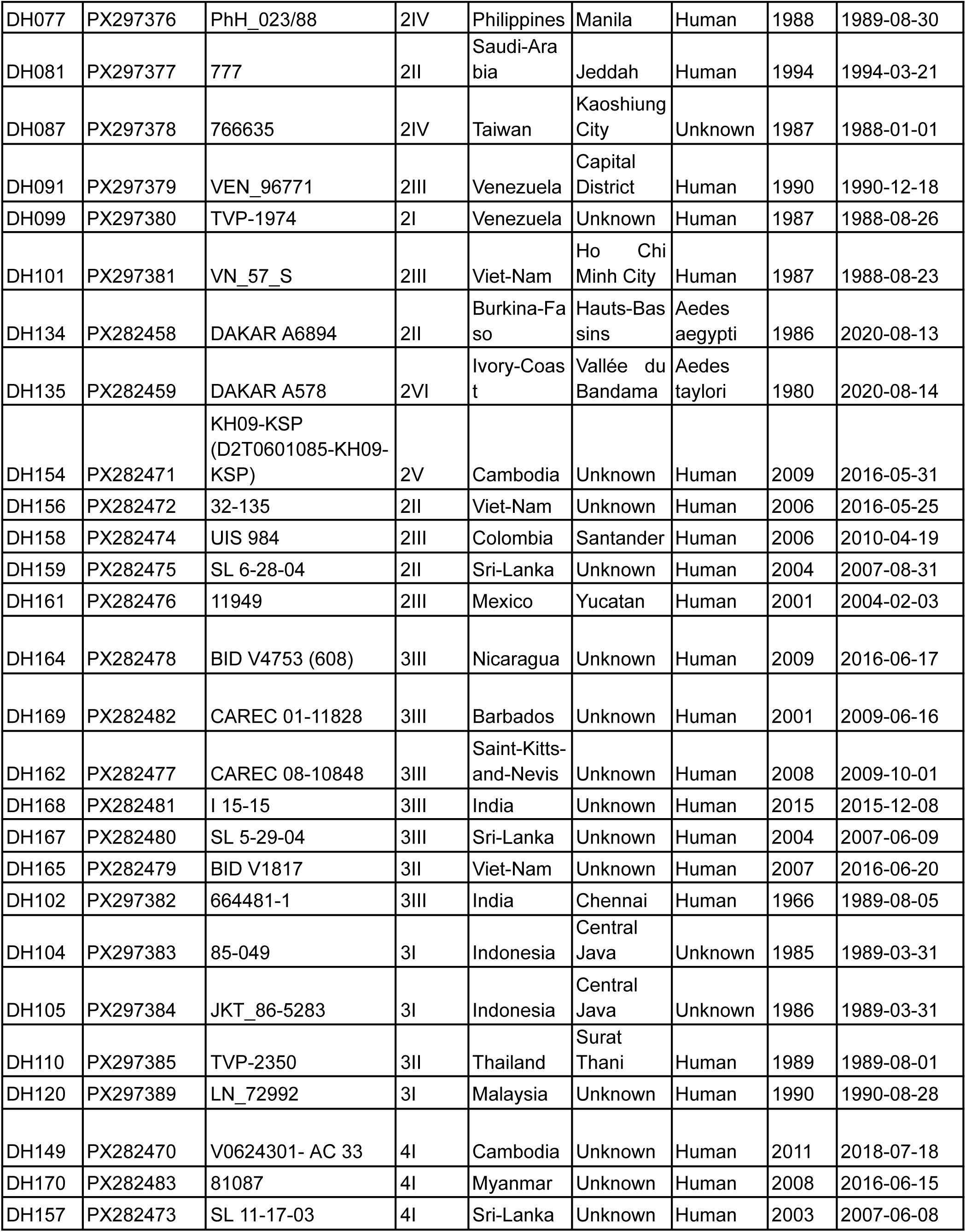
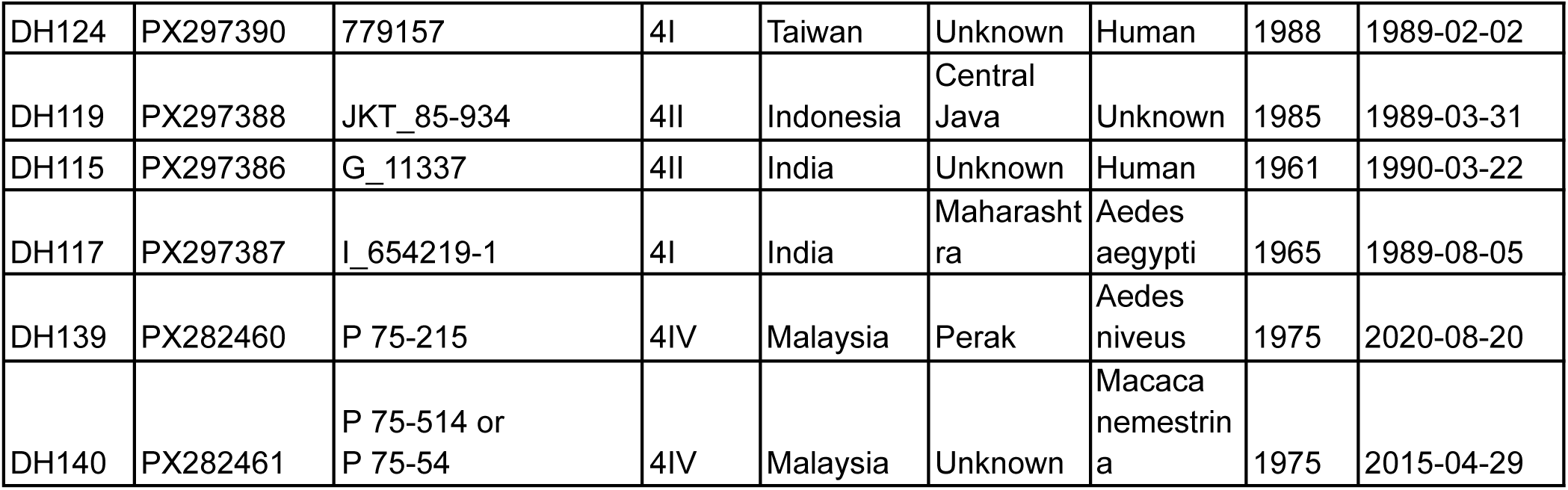
Metadata for dengue virus (DENV) strains included in this study. We received dengue virus isolates from the Yale Arbovirus Research Unit and World Reference Center for Emerging Viruses and Arboviruses. Our panel included 60 genetically diverse DENV strains spanning all four serotypes and the majority of genotypes. Each strain was characterized using metagenomic sequencing and submitted to NCBI GenBank under BioProject PRJNA1308108.

**Supplementary Table 2.**
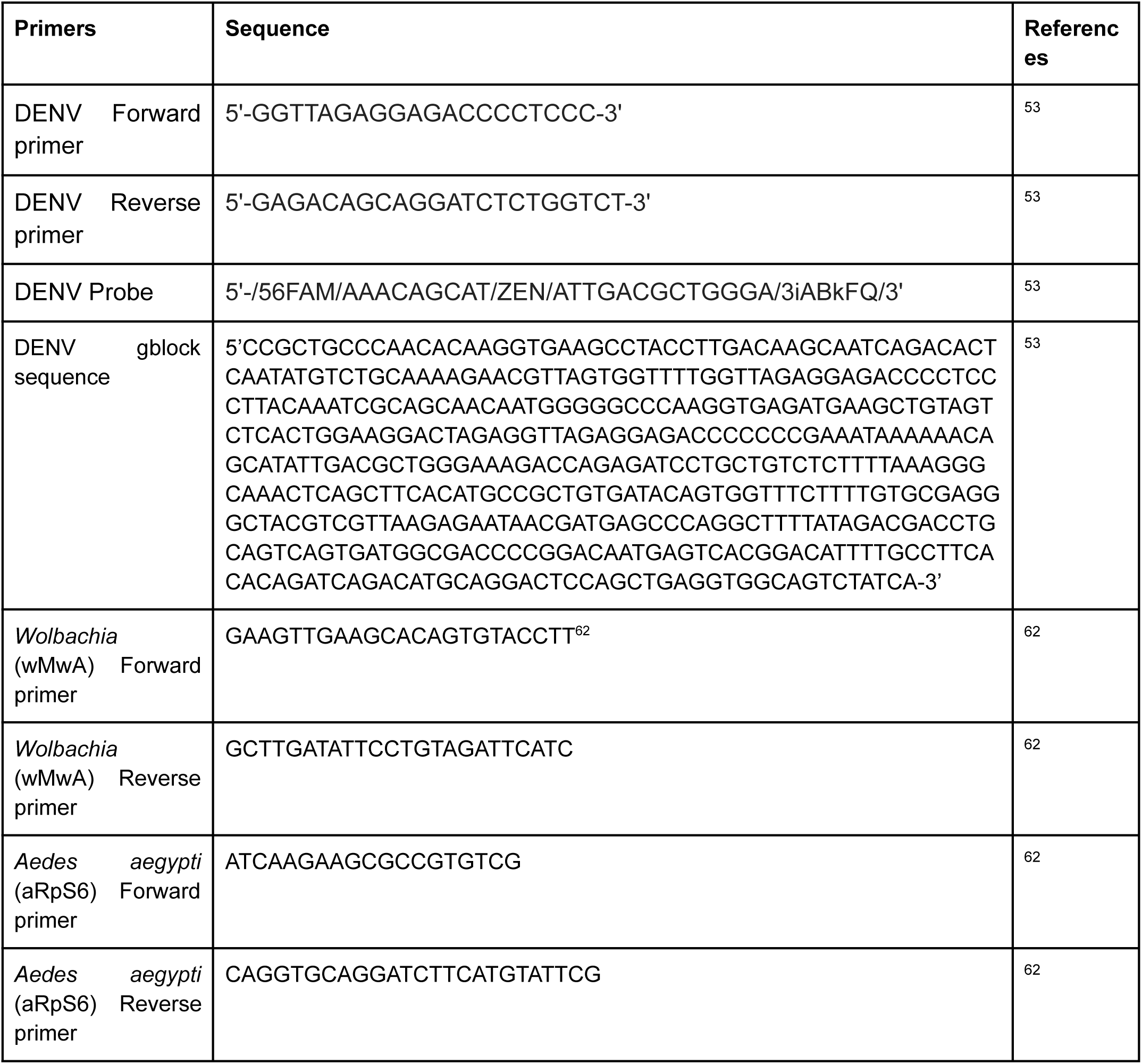
Primer, probe, and gblock sequences used for detection of dengue virus (DENV), *Wolbachia* (*w*AlbB and *w*MelM strains), and *Aedes aegypti*.

**Supplementary Table 3.** Fixed model parameters.

| Parameter | Description | Value | Reference/notes |
| --- | --- | --- | --- |
| $R_0$ | Basic reproduction number | 2-5 | Imai et al 2015<sup>63</sup> |
| $\gamma$ | Recovery rate | 0.2 | Duong et al 2015, Gubler et al 1981, Murgue et al 2000<sup>64–66</sup> |
| $1/\nu$ | Duration of cross-protection | 1 year | Reich et al 2013<sup>67</sup> |
| $\beta_1$ | Amplitude of cosine seasonality function | 0.2 | Assumed |
| $\varphi$ | Phase shift of cosine function | 1.56 | Based on exploratory analysis of Brazil's dengue seasonality<sup>68</sup> |
| $\lambda_{ext}$ | External FOI (representing introductions) | 1e-8 | Assumed |

## References

1. CDC. Areas with Risk of Dengue. Dengue https://www.cdc.gov/dengue/areas-with-risk/index.html (2025).

2. Engelthaler, D. M., Fink, T. M., Levy, C. E. & Leslie, M. J. The Reemergence of Aedes aegypti in Arizona - Volume 3, Number 2—June 1997 - Emerging Infectious Diseases journal - CDC. doi:10.3201/eid0302.970223.

3. 3. Dengue - Global situation. https://www.who.int/emergencies/disease-outbreak-news/item/2024-DON518.

4. Haider, N. et al. Global Dengue Epidemic Worsens with Record 14 Million Cases and 9,000 Deaths Reported in 2024. Int. J. Infect. Dis. IJID Off. Publ. Int. Soc. Infect. Dis. 107940 (2025) doi:10.1016/j.ijid.2025.107940.

5. Barcellos, C., Matos, V., Lana, R. M. & Lowe, R. Climate change, thermal anomalies, and the recent progression of dengue in Brazil. Sci. Rep. 14, 5948 (2024).

6. Bhatt, S. et al. The global distribution and burden of dengue. Nature 496, 504–507 (2013).

7. Childs, M. L., Lyberger, K., Harris, M., Burke, M. & Mordecai, E. A. Climate warming is expanding dengue burden in the Americas and Asia. medRxiv 2024.01.08.24301015 (2025) doi:10.1101/2024.01.08.24301015.

8. Moreira, L. A. et al. A Wolbachia Symbiont in Aedes aegypti Limits Infection with Dengue, Chikungunya, and Plasmodium. Cell 139, 1268–1278 (2009).

9. Ryan, P. A. et al. Establishment of wMel Wolbachia in Aedes aegypti mosquitoes and reduction of local dengue transmission in Cairns and surrounding locations in northern Queensland, Australia. Gates Open Res. 3, 1547 (2020).

10. Utarini, A. et al. Efficacy of Wolbachia-Infected Mosquito Deployments for the Control of Dengue. N. Engl. J. Med. 384, 2177–2186 (2021).

11. Lim, J. T. et al. Efficacy of Wolbachia-mediated sterility to reduce the incidence of dengue: a synthetic control study in Singapore. Lancet Microbe 5, e422–e432 (2024).

12. Nazni, W. A. et al. Establishment of Wolbachia Strain wAlbB in Malaysian Populations of Aedes aegypti for Dengue Control. Curr. Biol. CB 29, 4241–4248.e5 (2019).

13. Edenborough, K. M., Flores, H. A., Simmons, C. P. & Fraser, J. E. Using Wolbachia to Eliminate Dengue: Will the Virus Fight Back? J. Virol. 95, e02203–20.

14. Thi Hue Kien, D., et al. Genome evolution of dengue virus serotype 1 under selection by Wolbachia pipientis in Aedes aegypti mosquitoes. Virus Evol. 9, vead016 (2023).

15. Dengue Viruses | Learn Science at Scitable. https://www.nature.com/scitable/topicpage/dengue-viruses-22400925/#.

16. Sim, S. & Hibberd, M. L. Genomic approaches for understanding dengue: insights from the virus, vector, and host. Genome Biol. 17, 38 (2016).

17. Joubert, D. A. et al. Establishment of a Wolbachia Superinfection in Aedes aegypti Mosquitoes as a Potential Approach for Future Resistance Management. PLoS Pathog. 12, e1005434 (2016).

18. Corrêa-Antônio, J. et al. DENV-1 Titer Impacts Viral Blocking in wMel Aedes aegypti with Brazilian Genetic Background. Viruses 16, 214 (2024).

19. Gu, X. et al. A wMel Wolbachia variant in Aedes aegypti from field-collected Drosophila melanogaster with increased phenotypic stability under heat stress. Environ. Microbiol. 24, 2119–2135 (2022).

20. Flores, H. A. et al. Multiple Wolbachia strains provide comparative levels of protection against dengue virus infection in Aedes aegypti. PLOS Pathog. 16, e1008433 (2020).

21. Salje, H. & Jiggins, F. M. Risks of releasing imperfect *Wolbachia* strains for arbovirus control. Lancet Microbe 5, 622–623 (2024).

22. Carpenter, A. & Clem, R. J. Factors Affecting Arbovirus Midgut Escape in Mosquitoes. Pathogens 12, 220 (2023).

23. Salazar, M. I., Richardson, J. H., Sánchez-Vargas, I., Olson, K. E. & Beaty, B. J. Dengue virus type 2: replication and tropisms in orally infected Aedes aegypti mosquitoes. BMC Microbiol. 7, 9 (2007).

24. Gloria-Soria, A., Brackney, D. E. & Armstrong, P. M. Saliva collection via capillary method may underestimate arboviral transmission by mosquitoes. Parasit. Vectors 15, 103 (2022).

25. Dabo, S., et al. Extensive variation and strain-specificity in dengue virus susceptibility among African Aedes aegypti populations. PLoS Negl. Trop. Dis. 18, e0011862 (2024).

26. Johnson, R. M. et al. Implications of successive blood feeding on Wolbachia-mediated dengue virus inhibition in Aedes aegypti mosquitoes. 2025.02.06.636928 Preprint at 10.1101/2025.02.06.636928 (2025).

27. Ross, P. A. et al. Developing Wolbachia-based disease interventions for an extreme environment. PLOS Pathog. 19, e1011117 (2023).

28. Miller, B. R., Beaty, B. J., Atiken, T. H. G., Eckels, K. H. & Russell, P. K. Dengue-2 Vaccine: Oral Infection, Transmission, and Lack of Evidence for Reversion in the Mosquito, Aedes aegypti. Am. J. Trop. Med. Hyg. 31, 1232–1237 (1982).

29. Richards, S. L., Pesko, K., Alto, B. W. & Mores, C. N. Reduced infection in mosquitoes exposed to blood meals containing previously frozen flaviviruses. Virus Res. 129, 224–227 (2007).

30. Novelo, M. et al. Intra-host growth kinetics of dengue virus in the mosquito Aedes aegypti. PLOS Pathog. 15, e1008218 (2019).

31. Hoffmann, A. A. et al. Successful establishment of Wolbachia in Aedes populations to suppress dengue transmission. Nature 476, 454–457 (2011).

32. Souza-Neto, J. A., Powell, J. R. & Bonizzoni, M. Aedes aegypti vector competence studies: A review. Infect. Genet. Evol. J. Mol. Epidemiol. Evol. Genet. Infect. Dis. 67, 191–209 (2019).

33. Ekwudu, O. et al. Effect of Serotype and Strain Diversity on Dengue Virus Replication in Australian Mosquito Vectors. Pathog. Basel Switz. 9, 668 (2020).

34. Mancini, M. V. et al. High Temperature Cycles Result in Maternal Transmission and Dengue Infection Differences Between Wolbachia Strains in Aedes aegypti. mBio 12, e00250–21 (2021).

35. Ross, P. A. et al. A wAlbB Wolbachia Transinfection Displays Stable Phenotypic Effects across Divergent Aedes aegypti Mosquito Backgrounds. Appl. Environ. Microbiol. 87, e01264–21 (2021).

36. Ross, P. A. et al. Wolbachia Infections in Aedes aegypti Differ Markedly in Their Response to Cyclical Heat Stress. PLoS Pathog. 13, e1006006 (2017).

37. Ulrich, J. N., Beier, J. C., Devine, G. J. & Hugo, L. E. Heat Sensitivity of wMel Wolbachia during Aedes aegypti Development. PLoS Negl. Trop. Dis. 10, e0004873 (2016).

38. Ye, Y. H., Carrasco, A. M., Dong, Y., Sgrò, C. M. & McGraw, E. A. The Effect of Temperature on Wolbachia-Mediated Dengue Virus Blocking in Aedes aegypti. Am. J. Trop. Med. Hyg. 94, 812–819 (2016).

39. Walker, T. et al. The wMel Wolbachia strain blocks dengue and invades caged Aedes aegypti populations. Nature 476, 450–453 (2011).

40. Carrington, L. B. et al. Field- and clinically derived estimates of Wolbachia-mediated blocking of dengue virus transmission potential in Aedes aegypti mosquitoes. Proc. Natl. Acad. Sci. U. S. A. 115, 361–366 (2018).

41. Ferguson, N. M. et al. Modeling the impact on virus transmission of Wolbachia-mediated blocking of dengue virus infection of Aedes aegypti. Sci. Transl. Med. 7, 279ra37 (2015).

42. Londono-Renteria, B., Grippin, C., Cardenas, J. C., Troupin, A. & Colpitts, T. M. Human C5a Protein Participates in the Mosquito Immune Response Against Dengue Virus. J. Med. Entomol. 53, 505–512 (2016).

43. Giraldo-Calderón, G. I. et al. Transcriptome of the Aedes aegypti Mosquito in Response to Human Complement Proteins. Int. J. Mol. Sci. 21, 6584 (2020).

44. Ribeiro Dos Santos, G., et al. Estimating the effect of the wMel release programme on the incidence of dengue and chikungunya in Rio de Janeiro, Brazil: a spatiotemporal modelling study. Lancet Infect. Dis. 22, 1587–1595 (2022).

45. Garcia, G. de A., et al. Matching the genetics of released and local Aedes aegypti populations is critical to assure Wolbachia invasion. PLoS Negl. Trop. Dis. 13, e0007023 (2019).

46. Ross, P. A., Turelli, M. & Hoffmann, A. A. Evolutionary Ecology of Wolbachia Releases for Disease Control. Annu. Rev. Genet. 53, 93–116 (2019).

47. Paz-Bailey, G., Jernigan, D. B., Laserson, K., Zielinski-Gutierrez, E. & Petersen, L. New solutions against the dengue global threat: opportunities for *Wolbachia* interventions. Int. J. Infect. Dis. 157, 107923 (2025).

48. Velez, I. D., et al. Large-scale releases and establishment of wMel Wolbachia in Aedes aegypti mosquitoes throughout the Cities of Bello, Medellín and Itagüí, Colombia. PLoS Negl. Trop. Dis. 17, e0011642 (2023).

49. Matranga, C. B. et al. Enhanced methods for unbiased deep sequencing of Lassa and Ebola RNA viruses from clinical and biological samples. Genome Biol. 15, 519 (2014).

50. Illumina Pipeline Details. CZ ID Help Center https://chanzuckerberg.zendesk.com/hc/en-us/articles/360034790554-Illumina-Pipeline-Details (2025).

51. Ross, P. A. et al. Developing Wolbachia-based disease interventions for an extreme environment. PLOS Pathog. 19, e1011117 (2023).

52. Ross, P. A. et al. Developing Wolbachia-based disease interventions for an extreme environment. PLOS Pathog. 19, e1011117 (2023).

53. de Curcio, J. S. et al. Detection of Mayaro virus in *Aedes aegypti* mosquitoes circulating in Goiânia-Goiás-Brazil. Microbes Infect. 24, 104948 (2022).

54. Vasilakis, N. et al. Mosquitoes Put the Brake on Arbovirus Evolution: Experimental Evolution Reveals Slower Mutation Accumulation in Mosquito Than Vertebrate Cells. PLOS Pathog. 5, e1000467 (2009).

55. Lau, M.-J., Hoffmann, A. A. & Endersby-Harshman, N. M. A diagnostic primer pair to distinguish between wMel and wAlbB Wolbachia infections. PLoS ONE 16, e0257781 (2021).

56. Ross, P. A. et al. A decade of stability for wMel Wolbachia in natural Aedes aegypti populations. PLOS Pathog. 18, e1010256 (2022).

57. Huddleston, J. et al. Augur: a bioinformatics toolkit for phylogenetic analyses of human pathogens. J. Open Source Softw. 6, 2906 (2021).

58. Hadfield, J. et al. Nextstrain: real-time tracking of pathogen evolution. Bioinformatics 34, 4121–4123 (2018).

59. Dudas, G. evogytis/baltic. (2025).

60. Reich, N. G. et al. Interactions between serotypes of dengue highlight epidemiological impact of cross-immunity. J. R. Soc. Interface 10, 20130414 (2013).

61. World Population Prospects. https://population.un.org/wpp/downloads?folder=Standard%20Projections&group=Most%20used.

62. Lee, S. F., White, V. L., Weeks, A. R., Hoffmann, A. A. & Endersby, N. M. High-Throughput PCR Assays To Monitor Wolbachia Infection in the Dengue Mosquito (Aedes aegypti) and Drosophila simulans. Appl. Environ. Microbiol. 78, 4740–4743 (2012).

63. Imai, N., Dorigatti, I., Cauchemez, S. & Ferguson, N. M. Estimating Dengue Transmission Intensity from Sero-Prevalence Surveys in Multiple Countries. PLoS Negl. Trop. Dis. 9, e0003719 (2015).

64. Murgue, B., Roche, C., Chungue, E. & Deparis, X. Prospective study of the duration and magnitude of viraemia in children hospitalised during the 1996–1997 dengue-2 outbreak in French Polynesia. J. Med. Virol. 60, 432–438 (2000).

65. Duong, V. et al. Asymptomatic humans transmit dengue virus to mosquitoes. Proc. Natl. Acad. Sci. U. S. A. 112, 14688–14693 (2015).

66. Gubler, D. J., Suharyono, W., Lubis, I., Eram, S. & Gunarso, S. Epidemic dengue 3 in central Java, associated with low viremia in man. Am. J. Trop. Med. Hyg. 30, 1094–1099 (1981).

67. Reich, N. G. et al. Interactions between serotypes of dengue highlight epidemiological impact of cross-immunity. J. R. Soc. Interface 10, 20130414 (2013).

68. Sinan Dengue/Chikungunya. https://portalsinan.saude.gov.br/sinan-dengue-chikungunya.

